# Computational Modelling of Nephron Progenitor Cell Movement and Aggregation during Kidney Organogenesis

**DOI:** 10.1101/2020.01.14.905711

**Authors:** Pauli Tikka, Moritz Mercker, Ilya Skovorodkin, Ulla Saarela, Seppo Vainio, Veli-Pekka Ronkainen, James P. Sluka, James A. Glazier, Anna Marciniak-Czochra, Franz Schaefer

**Affiliations:** Division of Pediatric Nephrology. Heidelberg University Center for Pediatrics and Adolescent Medicine, Heidelberg, Germany; Faculty of Biochemistry and Molecular Medicine, Biocenter Oulu, University of Oulu, Oulu, Finland; Institute of Applied Mathematics (IAM) and Interdisciplinary Center of Scientific Computing (IWR), Mathematikon, Heidelberg University, Germany; Department of Intelligent Systems Engineering and Biocomplexity Institute, Indiana University, Bloomington, Indiana, USA

**Keywords:** Early Nephrogenesis, CompuCell3D, Cellular Potts Model, Particle Swarm Optimisation, Self-Organizing Maps

## Abstract

During early kidney organogenesis, nephron progenitor (NP) cells move from the tip to the corner region of the ureteric bud (UB) branches in order to form the pretubular aggregate, the early structure giving rise to nephron formation. Chemotaxis and cell-cell adhesion differences are believed to drive cell patterning during this critical period of organogenesis, but the spatiotemporal organization of this process is incompletely understood.

We applied a Cellular Potts model to explore to how these processes contribute to directed cell movement and aggregation. Model parameters were estimated based on fitting to experimental data obtained in *ex vivo* kidney explant and dissociation-reaggregation organoid culture studies.

Our simulations indicated that optimal enrichment and aggregation of NP cells in the UB corner niche requires chemoattractant secretion from both the UB epithelial cells and the NP cells themselves, as well as differences in cell-cell adhesion energies. Furthermore, NP cells were observed, both experimentally and by modelling, to move at higher speed in the UB corner as compared to the tip region where they originated. The existence of different cell speed domains along the UB was confirmed using self-organizing map analysis.

In summary, we demonstrated the suitability of a Cellular Potts Model approach to simulate cell movement and patterning during early nephrogenesis. Further refinement of the model should allow us to recapitulate the effects of developmental changes of cell phenotypes and molecular crosstalk during organ development.

**Author Summary:** The emergence of tissue patterns during vertebrate development is a major interest of both experimental research and biocomputational modelling. In this study, we established a Cellular Potts Model to explore cellular processes during early kidney development. The goal was to elucidate movements and aggregations of nephron progenitor cells. These precursor cells derive from mesenchymal cells around the ureteric buds and eventually form the epithelial structure of the nephron. Moreover, we wanted to explore computationally the mechanisms how these cells segregate from metanephric mesenchyme and move towards the location where the nephron will be formed. Utilizing the Compucell3D simulation software, we developed a model which assumes that nephron progenitor movement and aggregation is governed by only two mechanisms, i.e. cell-cell adhesion differences between cell types and nephron progenitor cell chemotaxis in response to chemoattractant secretion from two sources. These sources were either the epithelial cells of a static ureteric bud and/or the nephron progenitor cells themselves. The simulations indicated faster average cell speeds near the ureteric bud corner, the target region of cell movement and aggregation, and slower speeds near the place of origin, the tip of ureteric bud. The results were validated by comparison of the model predictions with experimental data from two ex vivo embryonic kidney models and a computational optimization protocol.

## Introduction

The mammalian kidney is the product of a highly complex, orchestrated developmental process, which involves not only proliferation and differentiation processes but also directed movement and aggregation of progenitor cells (Krause *et al*., 2015). Early nephrogenesis is characterized by the interplay of the branching and expanding ureteric bud (UB), the epithelial precursor structure destined to become the urinary tract, and the ‘cap’ metanephric mesenchyme (CM) surrounding the tips of the UB branches (Fig. 1) (BioPortal, 2019; Blake and Rosenblum, 2014; Bohnenpoll and Kispert, 2014; Costantini and Kopan, 2010; Desgrange and Cereghini, 2015; Obara-Ishihara T, 1999). A fraction of CM cells differentiate into nephron progenitor (NP) cells, which migrate towards the corner of the UB branches where they condensate to form circular pretubular aggregates (PTA) (Little, 2012). The PTA transforms into the renal vesicle, from which the final structures of the nephron, i.e. the tubules and glomeruli, are derived in a process of elongation and invagination (BioPortal, 2019; Combes *et al*., 2016).

**Fig. 1.**
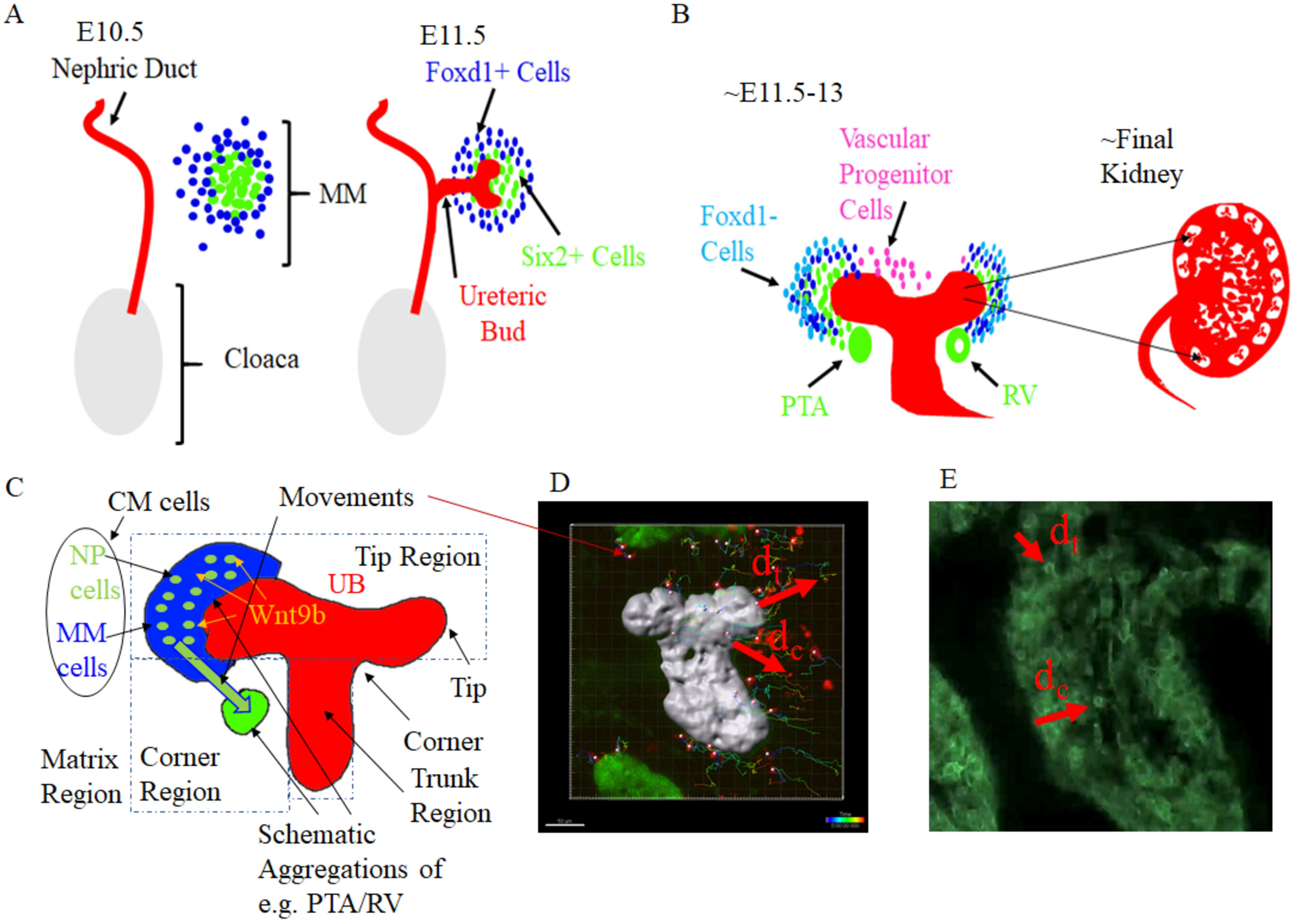
Morphological structures in early nephrogenesis. A) First branching of ureteric bud (UB) surrounded by cells of metanephric mesenchyme (MM). B) Terminal branch of UB with cap mesenchyme (CM), pre-tubular aggregate (PTA) and renal vesicle (RV). The places of UBs are shown in relation to a schematic figure of final kidney. C) Schematic drawing of UB trunk, corner, and tip regions with CM of NP and MM cells, with targeted movement of NP cells. D) Tracks of NP cell (red) movements around UB (grey) in kidney explant culture model. Scalebar (bottom left) represents 50 µm, c.f., (Combes et al., 2016). E) Kidney organoid model with black areas representing UB, and GFP labelled MM cells. D/E) Red arrows represent vectors of tip distance (dt) and corner distance (dc) used for analysis. The figure D was by a courtesy of Dr Alexander N. Combes regarding the similar figure in (Combes et al., 2016).

Early nephrogenesis research presumed that the movement of NPs towards the PTA was mostly linear (Little, 2012; McMahon, 2016). Recent studies indicate that NP movements are semi-stochastic and swarming-like (Combes *et al*., 2016), driven by adhesion differences and/or chemotaxis (Chen *et al*., 2015; Combes *et al*., 2016). The molecular mechanisms driving NP cell induction and PTA formation have been partially unravelled (Chen *et al*., 2015; Chi P *et al*., 2009; Combes *et al*., 2016; Little, 2015; Little, 2012). UB epithelial cells secrete various diffusible signalling proteins that may trigger the differentiation of MM to NP cells as well as their chemotactic movement towards the UB corner region (Combes *et al*., 2016; Little, 2015; Saarela *et al*., 2017). Furthermore, cell aggregation appears to be driven by differences in cell-cell adhesion properties (Lefevre *et al*., 2017), which may also be driven by autocrine and/or paracrine intercellular signalling (Dahl *et al*., 2002; Dudley *et al*., 1999; Oxburgh *et al*., 2011; Wallner *et al*., 1998).

A descriptive analysis of these processes is possible only to a limited extent, since even simple cell-cell interactions can lead to complex and unexpected large-scale spatiotemporal patterns (Magno *et al*., 2015). Therefore, computational methods have been introduced to mechanistically analyse early nephrogenesis (Combes *et al*., 2016). Combes et al., applying a convection-diffusion model, showed that attractive and repulsive cues between CM cells and the UB, together with cell adhesion processes, can lead to the commitment and maintenance of the CM in proximity to the tip (c.f., Fig. 1). However, the underlying cellular processes leading to the observed attraction and repulsion have not been analysed in detail, and the study was focused on the formation and maintenance of the CM rather than the formation of the PTA.

In this study we used a computational modelling approach to explore in detail the biophysical mechanisms underlying the directed movement and aggregation of NP cells, the critical first step of nephron development (BioPortal, 2019; Little, 2012). We extended the pioneering work of Combes et al. by systematically simulating chemical and mechanical cellular processes that potentially explain pattern formation during early nephrogenesis and used 3D tissue simulation approaches to analyse how different types of chemotaxis and adhesion differences between different cell types may explain the formation of both the CM and the PTA. Parameter calibration and model validation was achieved by comparison of the simulation results with both published and original experimental data.

## Results

### Initial Model Parameter Estimation

The initial parameter settings for *λ*_*A*_, *λ*_*V*_, *V*_*t*_, *A*_*t*_, and *J* (see Eq. 2 in Supplement) and the predefined spatial relationships defined the reference model (model 1, Table 1, Fig. 2A). Starting from these settings, the contact energy coefficients (*J*) between NP, MM, UB cells and the matrix were modified, yielding the adhesion-based models 2 and 8 (Table 1). The purpose of this modification was to identify the range of contact energy coefficients required to induce cell aggregations, and to explore the aggregation behaviour of NP and MM cells in the CC3D model space (Figs 2B/C) (Andasari *et al*., 2012; Combes *et al*., 2016; Osborne *et al*., 2017; Swat *et al*., 2009).

**Fig. 2.**
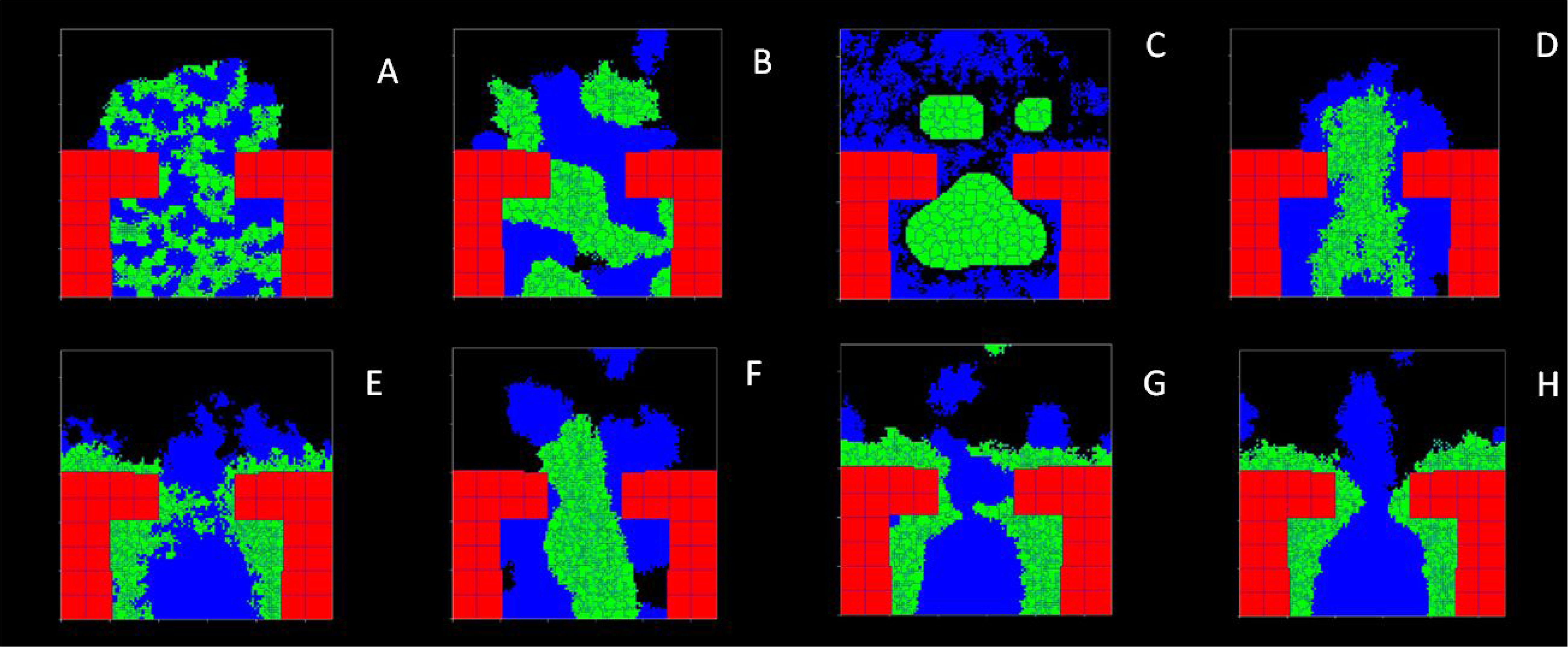
Final cell configurations obtained by 2D simulation with different models. A) 1_REF. B) 2_ADH. C) 8_ADH_ADH. D) 4_NP. E) 3_UB. F) 6_NP_ADH. G) 5_UB_ADH. H) 7_UB_NP_ADH. UB cells are depicted with red, NP cells with green, and MM cells with blue colours. Optimized parameters were calibrated to experimental cell properties and used in the simulations.

**Table 1.** Model parameters with pre-set values applied in 3D simulations.

| Parameter | Pre-set value | Models | Ref. |
| --- | --- | --- | --- |
| Target volume ($V_t$) | $375 \times 10^{-18} \text{ m}^3$ | 1-8 | this work,<br>(Combes<br>*et al.*,<br>2016) |
| Lambda volume ($\lambda_V$) | $20 \times 10^9 \text{ kgm}^{-4}\text{s}^{-2}$ | 1-8 | (Osborne<br>*et al.*,<br>2017;<br>Swat *et al.*, 2009) |
| Target surface area ($A_t$) | $312 \times 10^{-12} \text{ m}^2$ | 1-8 | this work,<br>(Combes<br>*et al.*,<br>2016) |
| Surface lambda ($\lambda_A$) | $0.1 \times 10^{-3} \text{ kgm}^{-2}\text{s}^{-2}$ | 1-8 | (Osborne<br>*et al.*,<br>2017;<br>Swat *et al.*, 2009) |
| Time step (MCS) | 0.017 h | 1-8 | (Osborne<br>*et al.*,<br>2017;<br>Swat *et al.*, 2009) |
| Surface temperature ($T$) | 20 | 1-8 | (Osborne<br>*et al.*,<br>2017;<br>Swat *et al.*, 2009) |
| Contact energy coefficient ($J$) | (all cells): $5 \times 10^{-15} \text{ kgs}^{-2}$<br>(NP&MM): $13 \times 10^{-15} \text{ kgs}^{-2}$ | 1,3-4;<br>2, 5-7, 8* | (Osborne<br>*et al.*,<br>2017;<br>Swat *et al.*, 2009) |
| Chemoattractant secretion ($S$) | 3 DU/s | 3-7 | (Osborne<br>*et al.*,<br>2017;<br>Swat *et al.*, 2009) |
| Chemotaxis lambda ($\lambda_{CL}$) | $100 \times 10^{-27} \text{ kgm}^2\text{s}^{-2}$ | 3-7 | (Osborne<br>*et al.*,<br>2017;<br>Swat *et al.*, 2009) |
| Global diffusion constant ($D$) | $1.0 \times 10^{-12} \text{ m}^2\text{s}^{-1}$ | 3-7 | (Brown,<br>2011; |
|  |  |  | Osborne<br>*et al.*,<br>2017;<br>Swat *et al.*, 2009) |
| Global decay constant ($\gamma$) | $1.0 \times 10^{-7}s$ | 3-7 | (Osborne<br>*et al.*,<br>2017;<br>Swat *et al.*, 2009) |

\*see Table 2 for description of multiple adhesion interfaces in model 8.

Next, the impact of chemotaxis on cell patterning was investigated, assuming secretion of chemoattractants by UB cells and/or NP cells themselves. To that end, parameters related to chemoattractant secretion (*S*), diffusion (*D*), degradation (*γ*), and chemotaxis strength (*λ*_*CL*_) were introduced (see Eqs 2, 3 in Supplement). To derive plausible value ranges for these parameters and explore the effect of chemotaxis on cell clustering, NP cells (model 4; Fig. 2D) or UB cells (model 3; Fig. 2E) were assumed as alternative sources of chemoattractant secretion. Finally, the combined effects of the contact energies and chemotaxis were considered in the remaining models (5,6,7; Figs 2F-H).

### Simulated Pattern Formation

The eight model variants yielded distinctly different final cell patterns (Fig. 2). The initial conditions and parameters had only a limited effect to the final cell patterns (Fig S1, S2). Therefore, the following description refers to the model patterns obtained with optimized simulations using random initial cell distribution and no pre-formed chemoattractant gradient.

The reference model yielded a cell pattern without coherent clustering over time (Fig. 2A). The models involving chemoattractant secretion by NP cells with and without cell adhesion differences resulted in cell aggregates between the UB tips without adherence to the UB surface (Figs 2D/F). In the models assuming adhesion differences between MM and NP cells but no chemotaxis, streak- or ball-like clusters emerged throughout the inter-UB area (Figs 2B/C). In the models involving UB cell chemoattractant secretion, NP cells aggregated along the UB surface, with preference to the corner regions (Figs 2E/G/H).

Directed migration and preferential aggregation of NP cells in the UB corner was observed with the models 3, 5, and 7 (Figs 2E/G/H), resembling the formation of PTA in the corner region during nephrogenesis (Fig. 2C). The most consistent formation of NP cell clusters resembling PTAs was observed with model 7, which involves chemoattractant secretion by both UB and NP cells and stronger adhesion between NP and MM cells relative to other models (Figs 2H, 3B/E/H). The video of the model (7) behaviour can be viewed online (Tikka, 2019d).

**Fig. 3.**
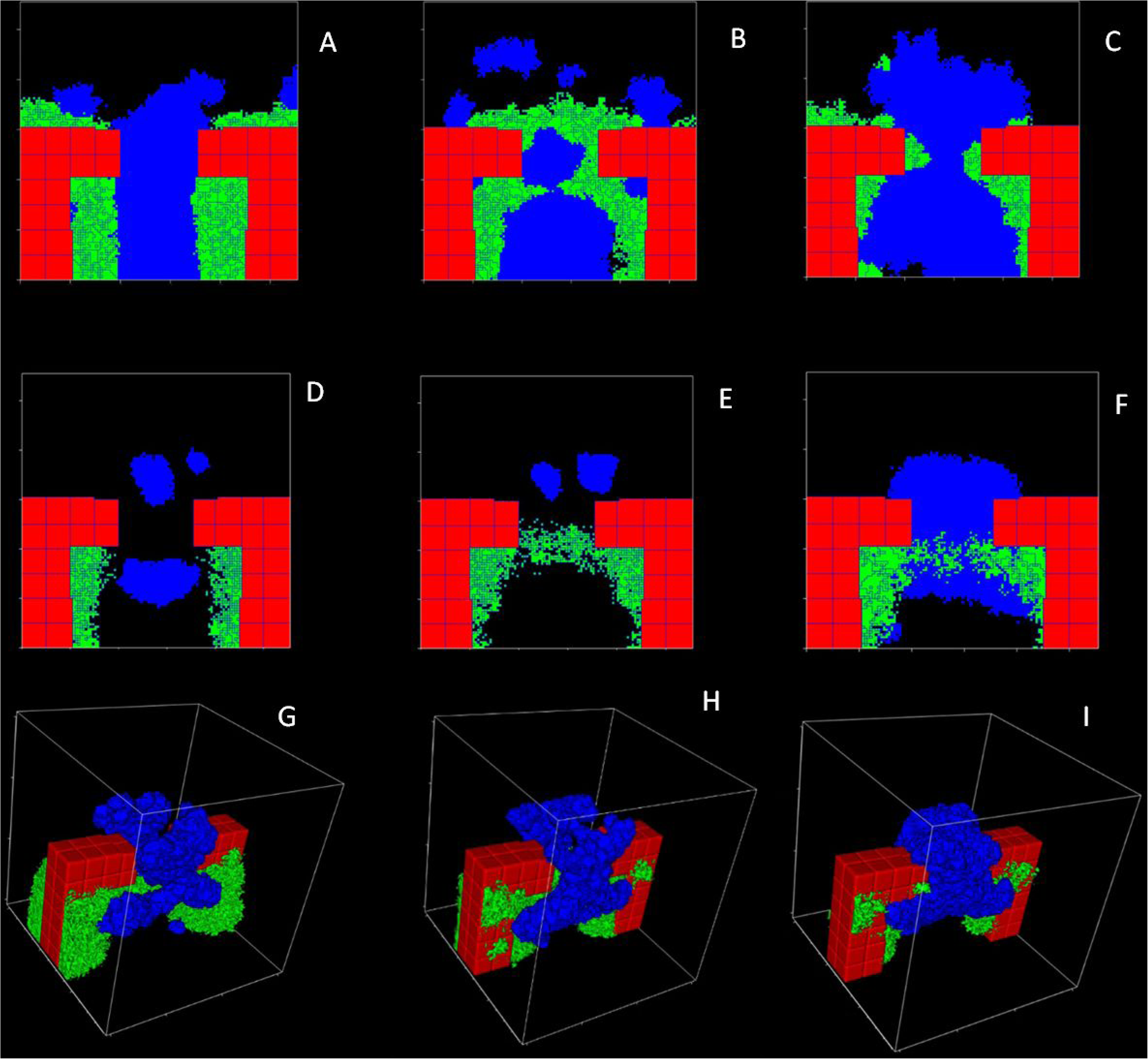
Representative final cell configurations obtained with the optimized models using initial chemoattractant fields. Model simulations of 5_UB_ADH_R (A/D/G) and 7_UB_NP_ ADH_R (A/D/G and B/E/H) were either in 2D (A-C) or 3D (D-I), assuming both the normal 50% or 25% (C/F/I) NP cells among total CM cell population. 3D views (G-I) are displayed together with their central transversal cuts (D-F). UB cells are depicted with red, NP cells with green, and MM cells with blue colours.

### Experimental Studies

The results of the cell movement analysis performed on the explant culture model experiments of Combes et al. (Combes *et al*., 2016; Lawlor *et al*., 2019; Lefevre *et al*., 2017) and our own experiments with a dissociation-reaggregation kidney organoid model are provided in (Figs 4-6).

**Fig. 4.**
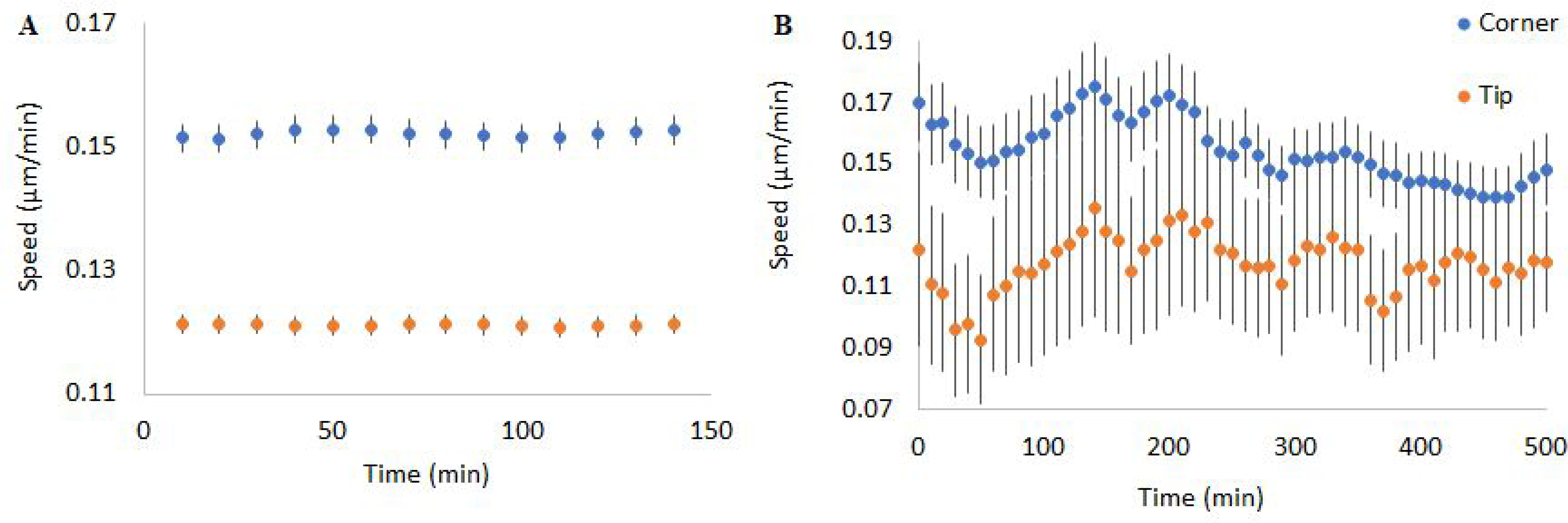
Cap mesenchyme cell speeds observed in (A) kidney organoid experiments, B) explant culture studies (Combes *et al*., 2016). Cell speeds in corner region are represented in blue and cell speeds around tip in orange. Dots represent means and vertical bars 95% confidence intervals.

The observed overall NP cell speed averages were 0.15±0.02 µm/min in the explant cultures, and 0.13±0.01 µm/min in the kidney organoid MM cells.

While cell speeds in the explant culture fluctuated considerably more than in the kidney organoid experiments, in both experimental settings the two cell types moved at different rates depending on their location relative to the UB tip (Fig. 4). In the corner region, the average cell speeds were 0.16±0.02 µm/min in the explant culture experiments and 0.15±0.01 µm/min in the kidney organoid (Combes *et al*., 2016). Average speeds in the tip region were 0.12±0.01 µm/min in both experimental settings. The slower relative cell movement of both MM and NP cells around the UB tip region was also apparent when expressed as the tip-to-corner speed ratio, which was below 1 for most of the observation time (Fig. 5).

**Fig. 5.**
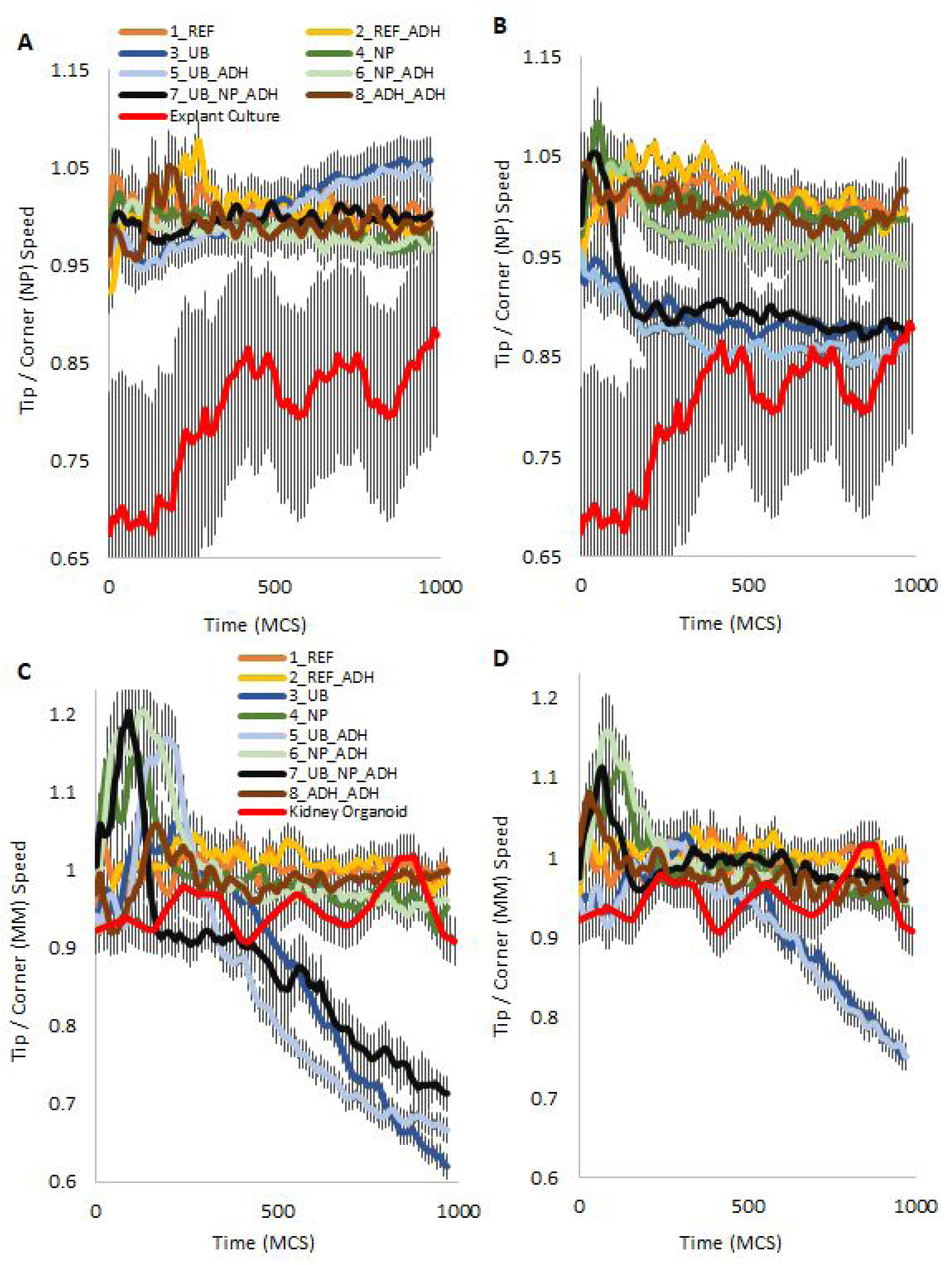
Average tip-to-corner speed ratios of NP and MM cells in the simulation studies of different models together with the results from experiments. A/B) NP cells in the models (see legend) with the cells in the explant culture experiments (scaled to model 3 and 4) (Combes *et al*., 2016). C/D) MM cells in the models similarly scaled with cells in the kidney organoid experiments. A/C) Simulations before the optimization. B/D) Simulations after the optimization. Vertical bars indicate 95% confidence intervals.

According to 2D SOM analysis, stable speeds in the explant culture data were 0.19±0.02 µm/min in the corner, and 0.14±0.01 µm/min in the tip region (Combes *et al*., 2016; Kohonen, 1982). 2D analysis was performed, because 3D data (z axis) of the explant culture experiments was found (regularly) skewed. Stable cell speeds of the kidney organoid data (calculated by 3D SOM analysis) were 0.25±0.02 µm/min in the corner and 0.18±0.02 µm/min in the tip region respectively. The different speed of motion of cells in the tip and corner regions is illustrated in the coloured speed contours of the transformed SOM plots (Fig. 6).

**Fig. 6.**
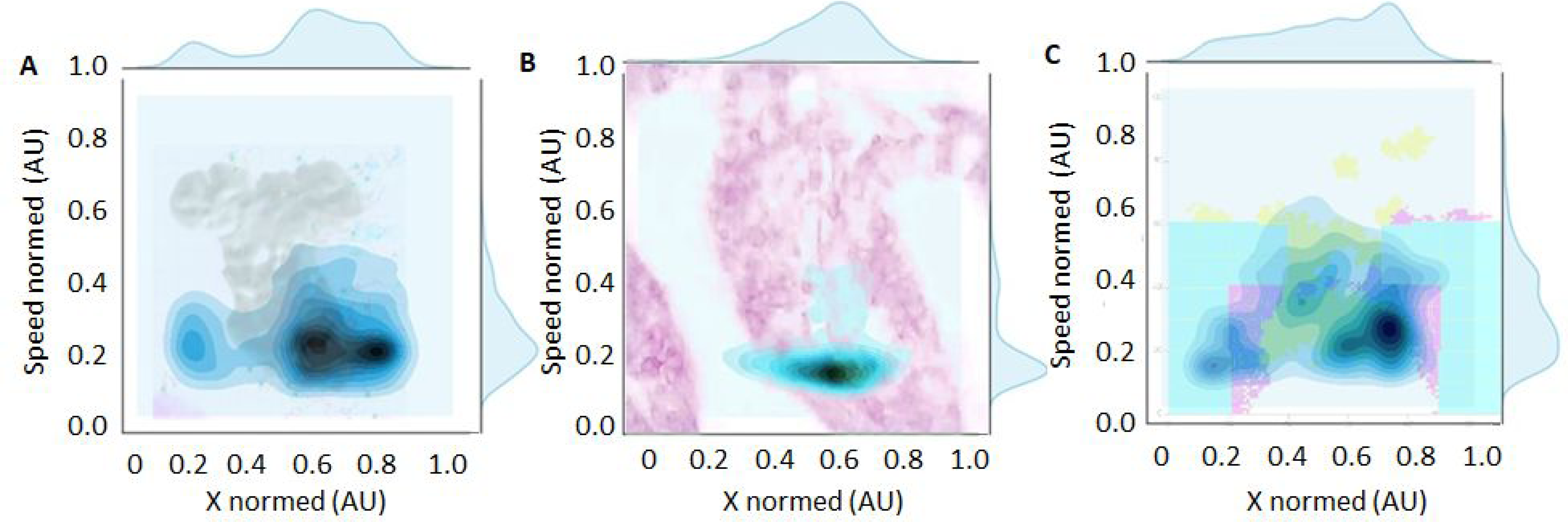
The cell speed contours of the best SOM groups (see supplement; (Kohonen, 1982)). A) NP cells in the explant kidney culture (Combes *et al*., 2016). B) MM cells in the kidney organoid model. C) NP cells in the optimized model seven. The speeds and the coordinates (x) have been normed (0,1). The experimental and simulation images in the background reflect the cell regions.

### Estimation of Final Model Parameters with Particle Swarm Optimization and Resulting Cell Patterns

The parameter ranges for PSO were chosen according to the initial model parameter estimates and the additional PSO algorithms (detailed further in the appendix). The optimization procedure aimed at maximising the amount of NP cells at the UB surface while simultaneously aligning the NP cell speeds in the model to the cell speeds observed experimentally in the explant culture setting, as mentioned in the methods.

The final model parameter values obtained by the PSO technique are given in Table 2. The improvement of the models achieved by the application of PSO is illustrated by the Best Quality Values (Table 2; lower numbers indicating better quality). The best model quality was obtained for model 7, while the other models showed either substantially lower quality or a spuriously high quality without matching the experimental situation. This applied in particular for the NP secreting models 4 and 6 and the adhesion model 8, where cells did not aggregate towards the UB surface in the first place.

**Table 2.** Optimized parameter values for each model variant (1-8). Values are presented as: Random (Uniform), e.g. for NP_ADH: 7.9 (13). {*} 8_ADH_ADH assumed nine cell-cell adhesion interfaces: {1-3} ‘Wall (UB) and NP/MM/Medium’, {4} ‘NP and NP’, {5} NP and MM’, {6} ‘MM and MM’, and {7-9} ‘Medium and NP/MM/Medium’. The parameters of spatial relationships (***V***_***t***_, ***A***_***t***_**, *λ***_***V***_, NO), except ***λ***_***S***_, and the ones not mentioned here were constant (see Table 1).

|  | REF<br>(1) | ADH<br>(2) | UB<br>(3) | UB_ADH<br>(4) | NP<br>(5) | NP_ADH<br>(6) | UB_NP_ADH<br>(7) | ADH_ADH<br>(8) |
| --- | --- | --- | --- | --- | --- | --- | --- | --- |
| Chemoattractant secretion rate (DU/s) | 0 | 0 | 3 | 3 | 3 | 3 | **6.91** | 0 |
| Chemotaxis lambda ($10^{-27}$; $\text{kgm}^2\text{s}^{-2}$) | 0 | 0 | **4.22**<br>(4.19) | **43.9** (43.8) | **67.94**<br>(71.3) | **74.5**<br>(74.3) | **1.41** | 0 |
| Global diffusion constant ($10^{-12}$; $\text{m}^2\text{s}^{-1}$) | 0 | 0 | 1 | 1 | 1 | 1 | **1.83** | 0 |
| Global decay constant (s) | 0 | 0 | $1 \times 10^{-7}$ | $1 \times 10^{-7}$ | 0.002 | 0.002 | **$4.69 \times 10^{-7}$** | 0 |
| Surface motility (Potts "temperature") (DU) | **10** | 25 | 25 | 25 | 25 | 25 | **49.8** | 25 |
| Surface lambda ($10^{-3}$; $\text{kgm}^{-2}\text{s}^{-2}$) | 0.01 | 0.01 | 0.01 | 0.01 | 0.01 | 0.01 | **8.11** | 0.01 |
| Adhesion differences between | 5.0 {5} | **7.9 {5}** | 5.0 {5} | **6.68 {5}**<br>(6.98 {5}) | 5.0 {5} | **7.9 {5}** (13 {5}) | **7.86 {5}** | **21.1 {1}, 25.2 {2}, 1.75 {3}, 0.50 {4},** |
| cell types<br>($10^{-15}$; $\text{kg s}^{-2}$) | | | | | | | | 33.3 {5}, 37.7 {6},<br>35.8 {7}, 29.7 {8},<br>1.57 {9}<br>(26.6 {1}, 47.8 {2},<br>29.5 {3}, 0.50 {4},<br>20.5 {5}, 0.52 {6},<br>24.1 {7}, 1.54 {8},<br>38.1 {9}) |
| PSO Best Quality<br>Value | -61,269 | -56,436 | -201,902 | -206,525 | -202,545 | -205,261 | -213,787 | -340,361 |

In the optimized models, the calculated total distance travelled by the NP cells during the simulation period was between 90-160 µm. The optimization procedure resulted in a more accentuated difference in corner and tip cell speeds in the models involving chemoattractant secretion by the UB (3, 5, 7 in Figs 5, S3). The different speed of motion of cells in the tip and corner regions is also illustrated for model 7 by the coloured speed contours of the transformed SOM plots (Fig. 6C). The tip-to-corner speed ratio of NP cells in these models decreased with time, aligning with the NP cell speed ratio observed towards the end of the explant culture experiments (Fig. 5B). By contrast, the MM cell speeds in the kidney organoid experiments were better matched by the optimized NP secreting models (4 and 6; Fig. 5D). In the secreting models 3-7 the overall speeds of the NP cells both in the tip and corner regions consistently exceeded those of the MM cells (0.13±0.03 v. 0.03±0.02 µm/min; Fig. 5). Correspondingly, NP cells in the kidney organoid experiments were enriched in the corner according to UB secreting models (3, 5, 7; Fig. 7B). On the other hand, MM cells in the explant culture experiments increased in the tip similarly as in the NP secreting models or vice versa in the corner (4, 6; Figs 7, S4).

**Fig. 7.**
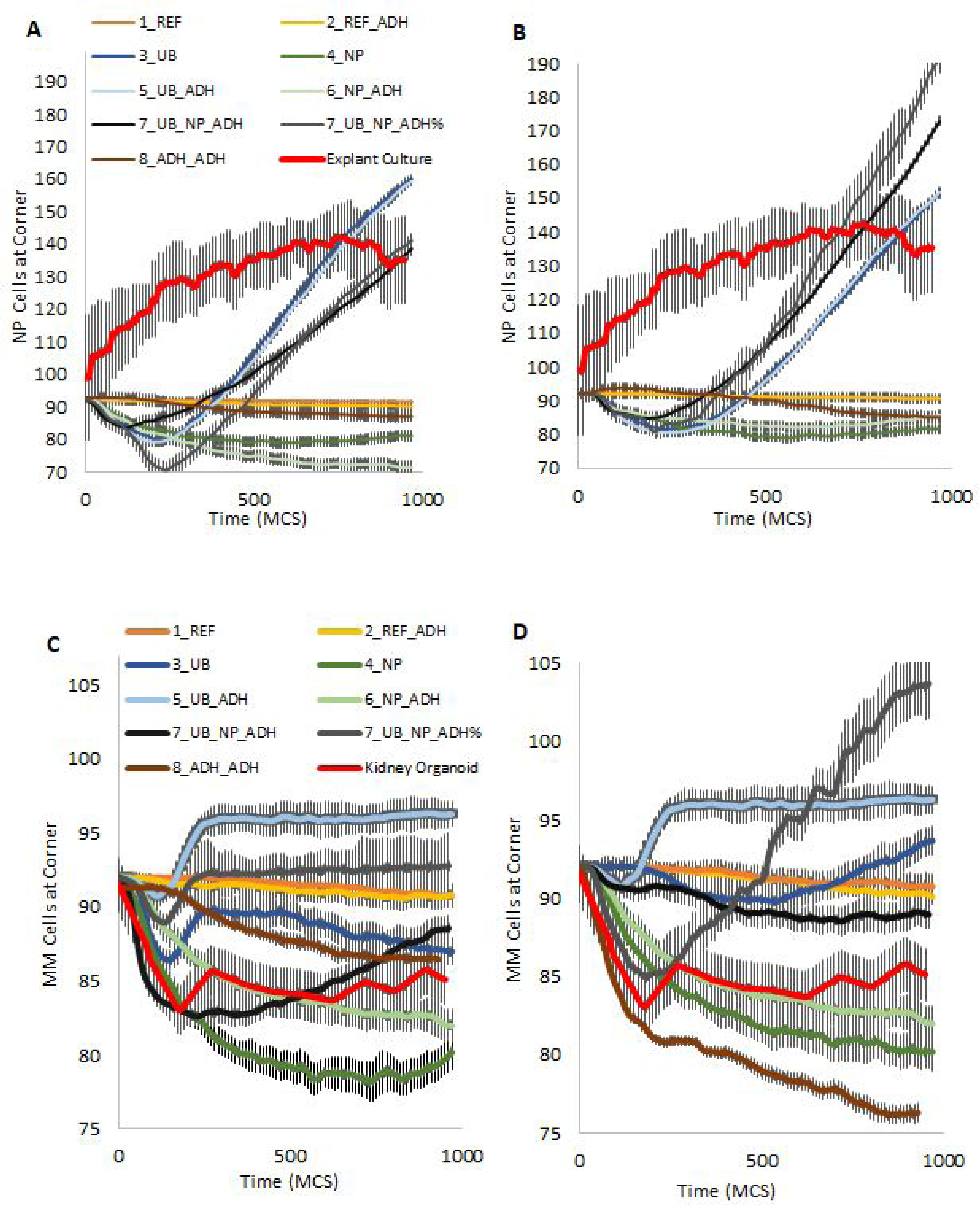
Enrichment of NP and MM cells in the corner region of the simulation studies of different models together with results from experiments. A/B) NP cells in the models (see legend) with the cells in the explant culture experiments (scaled to model 3 and 4) (Combes *et al*., 2016). Model 7 is presented with NP cell quantities of 50% (black; regular) and 25% (grey; scaled). C/D) MM cells in the similar models with cells in the kidney organoid experiments (see legend). A/C) Simulations before the optimization. B/D) Simulations after the optimization. Vertical bars indicate 95% confidence intervals.

The enrichment of the cells in different compartments is partly reflected by the relative tipper-corner cell distances of around 0.66 in the explant culture experiments, which were closest to the UB secreting models (3, 7; Figs S4, S5) (Combes *et al*., 2016). However, the relative tip-per-corner cell distances in the kidney organoids were best recapitulated by the adhesion-based or NP secreting models (1, 2, 4, 6; Fig. S5).

## Discussion

In this work we used a computational modelling approach to explore the biophysical mechanisms driving committed nephron progenitor cells to form proximal tubular aggregates, the early nephron precursor structure during kidney organogenesis. The movements and aggregations of the cap mesenchyme cells were studied using a Cellular Potts model, which was modulated to simulate the relative impact of chemotaxis and cell-cell adhesion forces. The movement of nephron progenitor cells towards the corner regions of the branching ureteric bud and the formation of cell aggregates was best reproduced by assuming a combination of chemotactic and differential cell adhesion forces. The parameter estimates were validated and optimized by analysis of cell behaviour in two experimental models of early nephrogenesis. We observed, both *ex vivo* and *in silico*, an accelerating speed of motion of committed NP cells as they migrate from the UB tip to the corner region.

Cap mesenchyme cells have been described to move in a quasi-stochastic fashion between the corner and tip regions of the branching ureteric buds (UB) following certain environmental cues (Combes *et al*., 2016; Lawlor *et al*., 2019). While the bulk of metanephric mesenchymal (MM) cells have been considered as static, an emerging subpopulation, the nephron progenitor (NP) cells, is believed to move linearly from the UB tip to the corner region (Little, 2012). In order to reproduce these movement patterns *in silico* several assumptions had to be made; these related to the initial spatial structure, cell quantities and properties and were founded on both experimental evidence and on established insights into cellular biophysics.

General assumptions included the assertion that all, and only, NP cells were committed to the formation of PTA, and that changes in three main energies should affect the movements and aggregations of NP and MM cells, i.e. contact surface energies, chemotaxis energy, and cell size changes. The impact of cell sizes was minimal since the initial most expansive phases were discounted in the simulation, leaving chemotaxis and cell-cell adhesion as the main energies to drive cell sorting between regions.

Indeed, there is substantial experimental evidence supporting the notion that the sorting of NP cells between the tip and corner regions is established by cell-cell adhesion differences and both inductive and chemotactic molecular signalling from the ureteric bud epithelia (Brown *et al*., 2013; Karner *et al*., 2011; Lefevre *et al*., 2017). Previous studies identified cell-cell adhesion molecules such as cadherins, to drive cell sorting (Junttila *et al*., 2015; Lefevre *et al*., 2017). Extracellular signals inducing NP cell commitment involve the secretion of WNT11, BMP7, FGF9, and WNT9B, which upregulates Wnt4 (Bohnenpoll and Kispert, 2014). NP cell induction has also been shown to involve the activation of Notch and additional signalling pathways (Lindstrom *et al*., 2015; Perantoni *et al*., 2005). At the same time, MM cells require SMAD1/5-mediated BMP signalling to transition towards a state in which they can receive the inductive cues (Brown *et al*., 2013). While WNT9B was the first secreted molecule attributed a role as a chemoattractant for NP cells, subsequent studies identified PDGF-AA, FGF8, BMP4 and CXCL12 as further potential effectors driving NP cell chemotaxis (Atsuta and Takahashi, 2015; Carroll *et al*., 2005; Grieshammer *et al*., 2005; Ricono, 2008). Recently, evidence has been provided that NP cells within the UB tip region do not move in a linear fashion but exhibit a nearly stochastic swarm-like behaviour (Combes *et al*., 2016). Moreover, NP cell commitment and migration towards the corner region may not be a unidirectional, irreversible process: a subset of NP cells at the corner region were observed to migrate back to the tip region to re-enter the uncommitted MM cell pool, losing Wnt4 expression (Lawlor *et al*., 2019). This behaviour was tentatively explained by semi-stochastic cell movement with exposure of NP cells to different cues depending on their spatial position, with prolonged or additional signals to NP cells being required for persistent clustering in the corner region and PTA formation.

We performed various model simulations to test the impact of chemotaxis and adhesion driven cell sorting on NP cell trafficking and clustering towards the region where PTA could be formed. The initial model setup and simulation studies were followed by a validation and optimization step, where explant culture and organoid experiments were utilized to further align the model parameters to fit the cell movements and aggregations observed *ex vivo*.

Our simulation studies support an important role of chemoattractant gradients arising from the UB surface for net directed NP cell movement. The model variants lacking UB chemoattractant secretion resulted in ectopic NP cell aggregation. Moreover, differential cell-cell adhesion properties appear to be required for the formation of NP cell aggregates. Paracrine chemotactic signalling by NP cells may play a role in this process as suggested by the optimal performance observed with a model combining adhesion differences with chemoattractant secretion by both the UB and the NP cells in the model 7.

Our model also adequately recapitulated the semi-stochastic movement of NP cells around the ureteric buds. Both the average cell speed and their net travelled distance fitted the speeds and distances observed in the explant culture system. Notably, with the model 7 we even observed individual cells returning from the corner to the tip region, in keeping with recent experimental findings (Lawlor *et al*., 2019).

Another notable finding, obtained both in the analysis of the explant culture data and by computational modelling and confirmed by self-organized speed mapping, is the faster NP cell speed in the corner as compared to the tip region. Our model simulations indicated that the acceleration of NP cells approaching the UB corner is due to the high chemoattractant gradient present in this region.

The behaviour of cells in the dissociation-reaggregation kidney organoid culture experiments was more challenging to simulate since no marker differentiating native MM from committed NP cells was available. As in the explant culture study, persistently faster cell movement was observed for cells in the corner and slower speed for those in the tip region. In contrast to the explant culture studies where our model simulations pointed to the UB as the predominant source of chemoattractant, regional cell speeds and quantities in the organoid studies were best approximated by the simulation assuming chemoattractant release from NP cells in addition to differences in cell-cell adhesion energies (model 6). This difference might have been caused by altered secretory functions of the ureteric bud epithelia following cell dissociation and reaggregation, and by the shorter experimental time period utilized in the kidney organoid model which was primarily designed to study early MM cell movement patterns. Nonetheless, the model 6 indicates that mesenchymal cell movement may be primarily driven by auto/paracrine chemotaxis of mesenchymal cells.

While our work demonstrates the suitability of a relatively simple computational model to reproduce the main cellular events in early nephrogenesis, several limitations should be emphasized. First, we did not allow for continuous recruitment of NP cells from the MM cell pool but assumed fixed cell quantities during the time window of analysis. Also, the lack of *in vitro* models deficient in individual components of the biological system prevented an external validation of the performance of our model in simulating impairments of nephrogenesis under abnormal conditions. Finally, the current model system did not allow to explore the roles of individual molecular signalling pathways or more than a single chemoattractant gradient. Such models would have required far more granular spatiotemporal biochemical information than currently available. Given these limitations, it is even more remarkable that NP cell migration and PTA formation can be accurately modelled based on two biophysical mechanisms, i.e. chemotaxis and cell-cell adhesion differences.

In conclusion, we established, validated, and applied a three-dimensional computational model of early nephrogenesis which describes the migration and initial aggregation of nephron progenitor cells as a function of stochastic swarming, chemotaxis and cell-specific adhesion properties. Our simulations suggest chemoattractant secretion by both the ureteric bud epithelia and the nephron progenitor cells themselves. Both experimentally and by simulation a non-linear migration pattern was observed, with cell movement accelerating while moving from the ureteric bud tip to the corner region. Our work demonstrates that computational modelling can aid in interpreting experimental data to reveal underlying biophysical mechanisms. The proposed model may be used as a starting point for more refined model systems as new molecular insights and experimental settings with higher information content become available.

## Methods

### Model Construction

We applied Cellular Potts model (CPM) to describe the behaviour (movement and aggregation) of NP and MM cells during early murine nephrogenesis (postconceptional day 12), and used the CompuCell3D software (CC3D) for this purpose (Combes *et al*., 2016; Lefevre *et al*., 2017; Saarela *et al*., 2017). Mathematical details of the developed CPM are given in the supplements (

Cellular Potts Model (CPM)). The following general assumptions were made to adjust CPM (Combes *et al*., 2016; Saarela *et al*., 2017):

1. NP, MM, and matrix cells are able to migrate by cell sorting (as described in CPM chapter) (Combes *et al*., 2016; Lawlor *et al*., 2019).
2. Matrix cells are chemically and physically neutral in comparison to other cells (Combes *et al*., 2016).
3. Only NP cells are able to perform chemotaxis, while UB, NP, or both cell types can secrete a chemoattractant (Junttila *et al*., 2015).
4. UB cells are static (“drift corrected UBs” (Combes *et al*., 2016)).
5. All cell types undergo neither proliferation nor apoptosis during the period of the observation (Combes *et al*., 2014; Lindström *et al*., 2018; Little, 2012).

The initial (3D) CC3D model setting comprised 2 L-shaped structures composed of 64 UB cells each separated by a space filled with MM and NP cells (n=196 each), and a matrix compartment comprising all remaining empty (pixel) space (Swat *et al*., 2009). The setting mimicked the spatial structure of two adjacent UB branches with surrounding metanephric mesenchyme (Fig. 8) (Andasari *et al*., 2012; Combes *et al*., 2016). The CM cells initially surrounding the UB tips consist of MM cells, which have similar cell volumes, masses, and general regulation mechanisms (

**Fig. 8.**
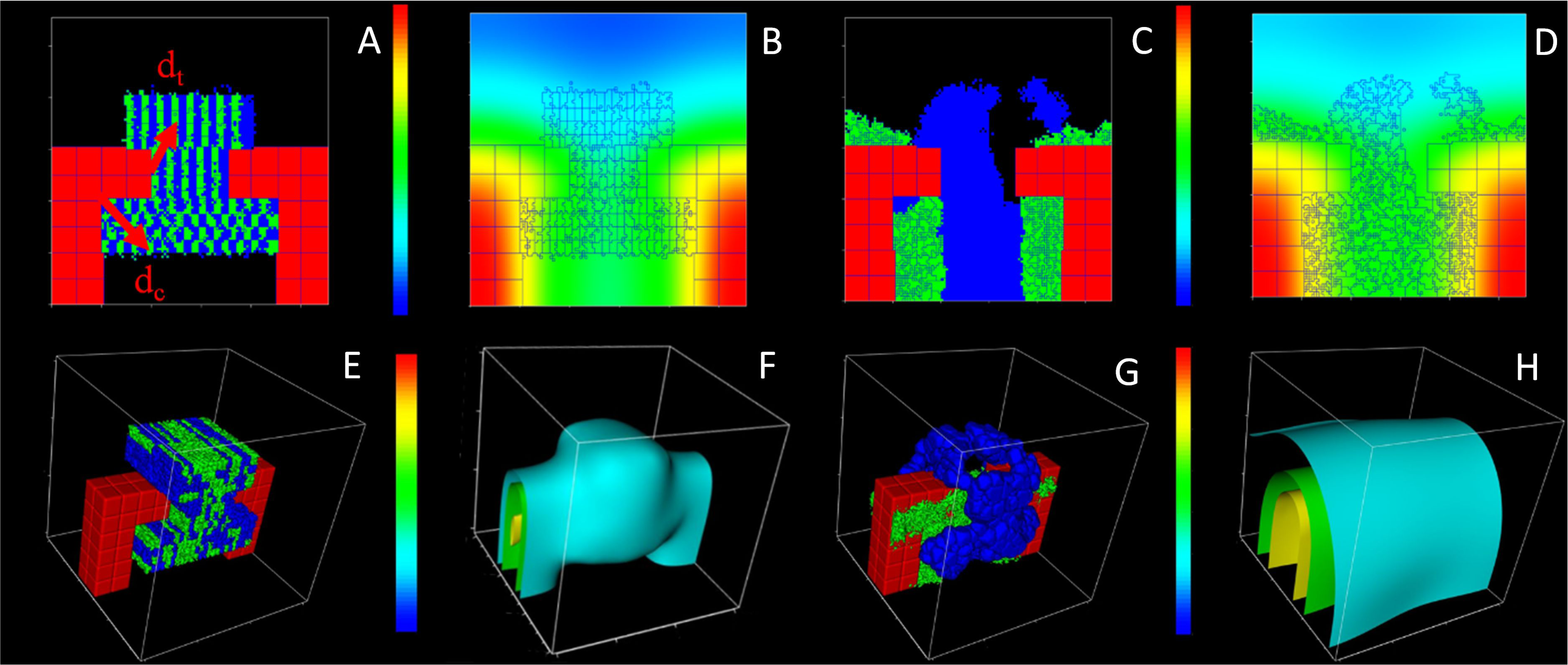
Two-dimensional (A-D) and three-dimensional (E-H) simulation patterns obtained with model 7_UB_NP_ADH. A/E) Initial cell patterns with uniform (A) and random (E) cell positions; tip distance (**d_t_**; [40,50]) and corner distance (**d_c_**; [20,40]) vectors are depicted in (A). B/F) Initial chemoattractant gradient patterns. C/G) Final cell patterns. D/H) Final chemoattractant patterns. In panels A, C, E, and G UB cells are depicted in red, NP cells in green and MM cells in blue, while matrix space appears black. In panels B, F, D, and H, standardized chemoattractant concentration gradients are depicted by coloured areas (2D) or sheets (3D), ranging from 0 (blue) to 1 (red).

Fig. 1C). NP cells are induced from CM cells at the outset of the modelled period (Lawlor *et al*., 2019). All cells were set to cubical shape, with initial cell surface areas of 375 µm^3^ for the MM and NP cells and 1000 µm^3^ for the UB cells. The distal end of the UB branch structure was denoted as “tip region” and the origin of the inner angle of UB as “corner region” (Fig. 8C). Our primary interest was to analyse the model cell outcomes in the corner region, where the PTA is formed (Fig. 8C-E).

### Simulation Studies

Eight model variants were constructed to simulate the impact of (a) chemotaxis of NP cells related to chemoattractant secretion from NP and/or UB cells separately or together with (b) adhesion-based cell sorting related to adhesion differences between different cell types. The model characteristics are listed in (Table 1). These variants represented the mechanisms potentially driving the cells in the studied experimental biological systems (*ex vivo* kidney and renal organoid), which corresponded the evaluated *in vivo* early nephrogenesis (Combes *et al*., 2016).

The first model variant (1_REF) did not include mechanical or chemical differences between different cell types but NP and MM cells showed patterns of ‘random walking’. This model served as a reference, or minimum model. The second model (2_REF_ADH) assumed differences in adhesion energies between NP and MM cells (Lefevre *et al*., 2017) but no chemotaxis processes. Models 3 and 4 introduced chemotaxis of NP cells, assuming chemoattractant secretion either by the UB cells (3_UB) or the NP cells themselves (4_NP), while adhesion properties of NP and MM cells were set equal. In models 5 and 6, models 3 and 4 were augmented by adding adhesion energy differences between NP and MM cells (5_UB_ADH, 6_NP_ADH). The seventh model included all features, i.e. chemotaxis of NP cells, adhesion differences between NP and MM cells, and chemoattractant secretion by both NP and UB cells (7_UB_NP_ADH). Finally, a multi-adhesion model (8_ADH_ADH) was tested with adhesion differences between all cells in the model (except UB-UB). For analysis and interpretation, models 3, 5, and 7 were categorized as ‘UB secreting models’, 4 and 6 as ‘NP secreting models’, 1, 2, and 8 as ‘non-secreting models’ and (2, 5, 6, 7, and 8) as ‘adhesion-based models’.

The following model characteristics were permutated in the models to test their effect on cell movement and aggregation in the different model variants: (i) Initial (NP and MM) cell positions; (ii) initial spread of the chemoattractant from UB or NP or both (Combes *et al*., 2016); (iii) cell-cell adhesion properties, especially of the moving NP and MM cells (Lefevre *et al*., 2017). For this purpose, we tested NP and MM cells with random (R) or uniform (U) initial cell distribution together with the spread of chemoattractant by simulations with or without the initial presence of a chemoattractant field (Fig. 8, Table 1).

Each model variant was simulated 30 times. During each of these basic simulations (8×30×4=960), we recorded both the centre of mass (COM) of the moving cell types (NP and MM) and the relative amount of chemoattractant in these COMs over 1000 Monte Carlo Steps (MCS, see Cellular Potts Model (CPM)). With these COM values, we calculated the average speeds, distances, and the local concentration of the chemoattractant separately in the tip and corner regions (Figs 1C, 8A, S7, see ‘Model Simulations’ and ‘Technical’). The cell speeds in the corner and tip regions were denoted as ‘corner speeds’ and ‘tip speeds’, respectively. Similarly, the distances of these cells from the outermost point of the UB tip were denoted as “tip distances” and the distances from the UB corner as “corner distances” (Figs 1C, 8A). For direct comparison, the ratio of the tip to the corner values of the speeds, distances, and concentrations were calculated. Exemplary 3D model simulation snapshots are shown in Fig. 8.

### Computational Estimation of Model Parameters

Two computational schemes were elaborated for estimating the parameters of the model variants in order to investigate the qualitative and quantitative differences in the respective simulations.

The process of model parameter estimation was initiated by varying values in a 2D cell-sorting simulation study using the CC3D software as described in the (back-inducing) strategy mentioned below (see also

Cellular Potts Model (CPM) in Appendix) (Andasari *et al*., 2012; Combes *et al*., 2016; Osborne *et al*., 2017; Swat *et al*., 2009; Swat *et al*., 2012). The initial parameters of spatial relationships (such as *λ*_*A*_, *λ*_*V*_, *V*_*t*_, *A*_*t*_ in Eq. 2) and the cell numbers were chosen to be in comparable ranges as observed in the explant culture and kidney organoid experiments (see experimental data sources and Figs 7, 8). These initial settings constituted the reference model 1. The initial parameter ranges for the contact energy coefficient (*J*), which refers to the cell-cell adhesion differences (Chen *et al*., 2015; Lefevre *et al*., 2017), and chemotaxis strengths (*λ*_*CL*_) (Chen *et al*., 2015; Chi P *et al*., 2009; Combes *et al*., 2016; Little, 2015; Little, 2012) were taken from the respective literature for the other 2D models (2-8) (Magno *et al*., 2015; Swat *et al*., 2009). The parameter values for all 2D models were subsequently scaled to 3D by multiplying by two as explained in (Magno *et al*., 2015), given the neighbourhood order of three. The final parameter values for the 3D models are given in Table 1.

The simulation strategy for the models 2-8 was to produce NP cell aggregates, preferably close to spheroidal shape, by permutating the previously mentioned initial parameters (Andasari *et al*., 2012; Lefevre *et al*., 2017; Saarela *et al*., 2017; Swat *et al*., 2009). Particularly, we aimed this aggregation to be directed towards the corner region at the end of the simulations (Figs 1C, 8A). Accordingly, the initial literature estimates were modified one at the time in the following order:

1. *λ*_*CL*_ of NP cells
2. *D*, *γ*, and *S* of the chemoattractant coming from UB and/or NP cells
3. *J* between NP, MM, and/or UB cells
4. *λ*_*V*_ and *λ*_*A*_ of NP and MM cells
5. *V*_*t*_ and *A*_*t*_ of NP and MM cells

The **Particle Swarm Optimization** (PSO) technique was used to optimize the model parameters (Anum *et al*., 2016). The PSO was set to maximise the number of NP cells at the surface of UB (in models 1, 2, 3, 5 and 8), mimicking the patterns in the kidney organoid experiments while simultaneously minimizing the difference between simulated and experimental (UB) tip cell speeds (Anum *et al*., 2016; Lefevre *et al*., 2017). In the NP secreting models (4, 6 and 7) we maximized the common surface area between NP cells for making PTA with the corner cell speeds. The construction of PSO codes for the models in the CC3D development program ‘Twedit++’ incorporated various model coding components presented in (Tikka, 2019a).

The optimization of the simulated cell speeds was achieved by comparison with the experimental data of Combes et al. (Combes *et al*., 2016) (see below), if the CM cells described in that study corresponded to the NP cells in our model. Simulation settings, parameters and constants that were optimized together using the PSO algorithm are referred to as ‘PSOed’. Details of the PSO method and the ranges of its parameters are given in the supplement.

Simulated cell speeds were also compared to the movement of MM cells observed in a kidney organoid model (see below), if these cells corresponded to both NP and MM cells in our simulations. Cells attached or close to the UB tip in the organoid model were considered MM cells in the simulations, while the remaining MM cells were assumed to be NP cells (7C-E, see also supplementary chapter ‘Technical Workflow’).

### Self-Organized Maps (SOM)

The Self-Organized Map (SOM) approach, an iterative machine learning method (Kohonen, 1982), was applied to identify and compare especially stable cell speed regions in the simulations and experiments. Specifically, we sought to compare if the simulated and real NP cells behaved differently at the tip and corner regions respectively. The Python function ‘MiniSom’ (GitHub, 2019) was applied to compare the SOM results derived from our simulations with the experimental data of Combes et al. (Combes *et al*., 2016). Details of the implementation are given in the supplement.

### Experimental Data Sources

In this study, two sources of *ex vivo* experimental data were used to calibrate and validate the simulation models (Combes *et al*., 2016; Tikka, 2019b). These comprised results obtained with an explant culture model reported by Combes et al. (2016), and original results reported in this work obtained with a dissociation-reaggregation kidney organoid model (Combes *et al*., 2016; Costantini and Kopan, 2010; Saarela *et al*., 2017; Tikka, 2019b). The data consisted of the images recorded during the experiments (see Microscopy methods below). The sizes of the recorded data frames (number of samples per number of time points) were 15 x 50 (for explant culture in 12.5-15h) and 1×15 (for kidney organoid study in 2.5h; see ‘Technical Workflow’ for more details). The explant culture experiments of Combes et al. comprised 500 NP cells, whereas the kidney organoid studies presented here comprised of 500-2000 MM cells. Outliers were replaced with a moving average of three previous data values; see python code ‘rolling mean’.

### Kidney Organoids

Embryonic kidneys were dissected from embryonic day 11.5 mouse embryos from crossing of Wnt4Cre (Shan *et al*., 2010) and tomato floxed Rosa26 Green fluorescent protein (GFP; mT/mG) reporter mice (Muzumdar *et al*., 2007) as described in (Junttila *et al*., 2015). Intact UBs were treated with GDNF and dissociated MM with BMP7 and FGF2 as in (Junttila *et al*., 2015). The intact UB was reaggregated with MM cells and incubated overnight to form a kidney organoid. The organoids were set to grow in a FiZD culture (Saarela *et al*., 2017) for time-lapse imaging into a temperature and gas controlled on-stage incubator (OkoLab, Italy) on a Zeiss LSM780 confocal microscope.

### Microscopy, Image Processing and Data Segmentation

In order to track and distinguish MM cells from UB cells, MM cells were expressing GFP. The fluorescence microscopy of this work was performed with a Zeiss LSM 780 confocal microscope (Carl Zeiss, Germany) using 25x/0.8 Zeiss LCI PlanNeofluar water immersion objective with a 488nm wavelength for excitation and a range of emission from 490-601nm. Out of original images of 250 µm x 300 µm x 29 µm (XYZ), an area of two close and relatively static UBs (approx. 150 µm x 100 µm) was selected with the Zen Blue (Carl Zeiss, Germany) imaging program. There were 16 Z-layers in the (z axis) stack from which all were selected. Here, XYZ pixel size was 0.24 µm x 0.24 µm x 1.46 µm and the temporal resolution was given by 16-time frames with 10 min between the frames. The video of the moving MM cells in the kidney organoid experiment is given stack-by-stack in (Tikka and Skovorodkin). Deconvolution was used to improve image signal to noise ratio, contrast and resolution. This process was done with Huygens Professional program (Scientific Volume Imaging, The Netherlands) using distilled PSF, with background value of 3.5, S/N value of 10, autobleach correction off, and saving the resulting deconvolved image as 16-bit tiff file format (Saarela *et al*., 2017). The segmentation of the tiff file of the deconvolved image was done with a previously compiled in-house MATLAB code (Saarela *et al*., 2017). The code produced a raw data file, .csv, of the positions, sizes and speeds of all of the cells by identifying objects that it redeemed as cells from the images. We specified the cell diameter range between 0.5-20 µm.

## Ethical Statement

Animal care and procedures were in accordance with Finnish national legislation for the use of laboratory animals, the European Convention for the protection of vertebrate animals used for experimental and other scientific purposes (ETS 123), and the EU Directive 86/609/EEC.

## Acknowledgements

PT received a grant from the Marie-Curie International Training Network RenalTract. JPS and JAG were supported by National Institute Health grants GM122424 and GM111243. FS is a member of the European Reference Network for Rare Kidney DIseases (ERKNet). We are grateful to Dr. Alexander L. Combes for allowing us to use the data frames obtained in their published explant culture experiments (Combes *et al*., 2016).

## Competing Interests

The authors declare no conflicts of interest.

## Author Contributions

PT performed the primary conceptual and programming work and drafted the manuscript. FS and MM helped writing the manuscript. MM helped in the model and analysis related questions. JAG and JPS assisted with developing the model in CompuCell3D and with the PSO simulations. IS and US conducted the kidney organoid experiments. VPS helped with the calibration of the microscopes and segmentation method. AMS and SV supported in the revisioning of the text. All authors approved the final version of the manuscript.

## Data Availability

The data of explant culture experiments have been published (Combes *et al*., 2016). The data of this work, including the experimental data in the kidney organoid and the analysis codes, have been uploaded to public repositories (Tikka, 2019a; Tikka, 2019b; Tikka, 2019c). The segmentation codes for the kidney organoid model are available at (Saarela *et al*., 2017).

## Supplemental Appendix

### Cellular Potts Model (CPM)

The CC3D models used here were based on the framework of CPM (Hirashima *et al*., 2017; Swat *et al*., 2012). During each simulated time step of the model, here named a Monte Carlo Step (MCS), all cell border pixels were attempted to be replaced by the neighbouring ones (Swat *et al*., 2012). This happened with a probability (*P*) according to Boltzmann distribution, which depended on the change in an ‘effective energy’ *H*

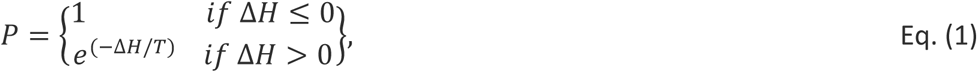

where *T* was called the temperature, i.e. the relative amount of cell surface random fluctuations. The main idea is that the displacements of individual cell pixels are accepted only if the overall effective energy (lower part of Eq. 2) is reduced. The power in the probability function depended also of the inverse of a “temperature”, which represents the effective motility of the cell’s membrane. If the net energy change after MCS was negative, i.e. Δ*H* ≤ 0, the index changes were adopted. In cases where Δ*H* > 0, the likelihood *P*, for a successful ‘index-copy attempt’, follows a Boltzmann distribution in (Eq. 1). The formulation of the effective energy contains quantities reflecting chemical and physical properties of the cells, such as area and volume constraints, cell-cell surface adhesion and processes related to chemotaxis (Andasari *et al*., 2012; Swat *et al*., 2012). Especially, the effective energy *H* represented a mix of true physical energies (such as cell-cell adhesion), terms that mimicked energies (e.g. the response of a cell to a chemotactic gradient) and terms that reflected basic principles of the evolution of the model (Hirashima *et al*., 2017). The effective energy used in this study is given by

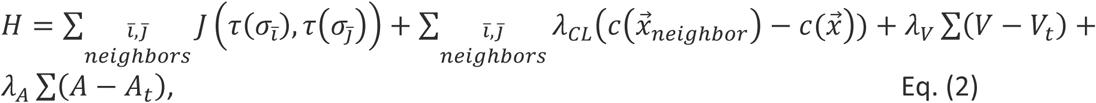

where the first sum denotes the adhesion energies, Δ*E*_*ad*ℎ*esion*_, and the second sum the chemotaxis energies, Δ*E*_*c*ℎ*emotaxis*_. Especially, *J* was defined as boundary energy per unit area between two different cells (*σ*_*i̅*_ *and σ*_*j̅*_) in the pixel interface of the cells ((*τ*(*σ*_*i̅*_), *τ*(*σ*_*j̅*_)). The chemotaxis energy term was calculated for chemotaxis cells and their neighbours. Here, *c* 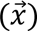 and *c*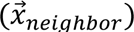 indicated the chemoattractant concentrations at the pixel source and its neighbouring pixel, respectively, where *λ*_*CL*_ reflected the strength of chemotaxis (Chen *et al*., 2015; Kopan, 2014; Swat *et al*., 2012). Finally, *λ*_*V*_ and *λ*_*A*_ penalized deviations of cell volume and area from the preferred values *A*_*t*_ and *V*_*t*_, respectively. A certain level of minimal adhesion (J) was necessary for the cell movements according to Eq. (2). The cells in a model without adhesion difference or chemoattractants would become nearly immobile soon after the beginning of the simulation. The volume expansion at the beginning of the simulation cannot sustain movements. Finally, the diffusion equation that governed the chemoattractant concentration evolution (*c*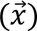) was

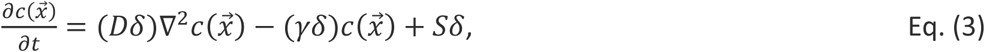

where *D*, *γ*, and *S* were the diffusion, degradation and secretion rates for the cell in question, such as NP, respectively. *δ* represent Boolean switches that were either 1 or 0 depending on whether the concentration was located on the cell or not. If it was in the cell in the question, then the switch value was one. The chemoattractant diffusion field 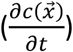 evolved independently of the cell domain in a separate domain (Fig. 8). However, this domain was similar in size as the cell domain (Fig. 8). The current modelling handled three cell types (UB, NP, MM) as the sources and receivers of the values of the field. Accordingly, the PDE in the equation (3) can be expanded into three equations (i.e. 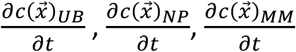) Similarly, all the constants and variables in each of the equations can be subscripted with the respective name of the cell, e.g. *D*_*NP*_ (Swat *et al*., 2012). ODEs could have been also used in respect to subcellular processes but were not considered in this modelling approach.

### Details of PSO Method

The search or iteration in PSO improves the candidate solutions with respect to a given measure of quality [15,28,31], such as the amount of NP cells in UB and speed differences. PSO solves the problem by using a population of candidate solutions. These solutions are called swarms. The algorithm moves the individual candidate solutions, i.e. particles, around the search-space according to the particle’s position and speed. The movement of particles are influenced by their own best-known positions. The movements are also guided towards the best-known position across all particles in the swarm. These best solution places are updated as better positions are found. This process moves the swarm towards the best solutions. PSO makes few assumptions about the problem being optimized and it can search very large parameter spaces efficiently (Andasari *et al*., 2012; Swat *et al*., 2009; Zhang *et al*., 2015).

The PSO parameters for the simulations were always the same. The momentum and scaling factors in equation 8 were *ω* =0.73 and *c*_1_=*c*_2_=1.5. The maximum speed for a parameter was limited to the parameter’s maximum allowed value. The PSO was implemented in python and the PSO code created the parameter files for each CC3D run, submitted the jobs to a SLURM job manager on a small Linux cluster, waited for job completion, extracted the results and calculated new parameters for each particle in each swarm.

The PSO’s were run for 60 iterations using 4 independent swarms and 8 particles per swarm. Since the CC3D simulations are stochastic, and replicate runs with the same parameter set do not give identical results, we ran each simulation three times and used the average result. Therefore, a complete PSO run for a model included 60×4×8×3=5760 simulations. For the 2D models a single CC3D simulation typically took 30 seconds, for 3D models 30 minutes. Since we had access to a 32-node cluster, we ran 32 simulations in parallel and the total time required for a model was therefore 60×3 times the time of a single run. Elapsed time for a complete PSO run was approximately 2 hours for the 2D models and 5 days for the 3D models.

As for the speed differences between the Combes et al. 2016 experiments and simulations, PSO penalized the overall quality value with difference of χ^2^ speed values in each (simulation and experiment) time point. This speed difference quality value was smaller than the quality value of the surface coverage (Anum *et al*., 2016; Combes *et al*., 2016). Accordingly, the speed quality values in each time point for every model were multiplied by scaling factor (of 100), to correspond to 20% of the maximum quality value from NP coverage at UB in 3D (7_UB_NP_ADH_R (3D)). It is unlikely that the speed quality values were higher in other models.

We applied the following constraints during the PSO iterations:

- Chemotaxis lambda (*λ*_*CL*_), secretion rate (*S*), the diffusion constant (*D*) as well as the decay constant (*γ*) were restricted by the upper bounds of 100, 30, 0.5 and 0.001, respectively; in order to allow an appropriate convergence of the CC3D algorithm.
- The initial contact energy coefficients (*J*) between NP and MM cell values in the adhesion increment models 2, 5, 6, 7, and 8 were 13 (vs. 5 *J* in models 1, 3, and 4).

Typical swarm parameter ranges were: *D* [0.1, 2.0], *γ* [10^-8^, 10^-6^], *J* [2.0, 8.0], *λ*_*CL*_ [10.0, 150.0], *S* [0.3, 30.0], *T* [5, 50.0], and Surface’s lambda [0.001, 10]. Depending on the model variant, PSOs yielded a variable number of PSOed parameters (Table 2).

7_UB_NP_ADH_R was deemed to be the most technologically advanced model in comparison to others. It contained all the values that other models had and more. Hence, it’s values could be used as a maximum appraisal that the other models should not exceed. For instance, the quality value in this model reached a plateau value of 220750. By comparison to other models, the quality values resulting from PSOs were in that order of magnitude. For example, 4_UB_R had a nominal value (in one PSO) of 199475, where the contribution of surface area was 186263.5. The contribution of the speed minimization in that model was a negative one (−13211.1). The set of optimized parameter values for this advanced model 7 were: *D*=1.83×10^-12^m^2^s^-1^ between (a range of 1, 2) with (PSO run time averages of 1.42±0.39) and *γ* =4.7×10^-7^×10^-7^s between (1×10^-8^, 1×10^-6^) with (7.2×10^-7^±1.7×10^-7^). It has also *J*=7.90×10^-15^ kgs^-2^ between (2, 8) with (6.49±2.23), *λ*_*CL*_ =14.1×10^-27^kgm^2^s^-2^ between (10, 150) with (15.2±5.1), *S*=6.91 DU/s between (0.3, 30) with (5.46±2.05), *T*=49.8 DU between (5, 50) with (48.14±1.15), and *λ*_*S*_=8.1×10^-3^kgm^-2^ s^-2^ between (0.001, 10) with (8.30±0.83). The 2D version of this model took 3.3h for 60 iterations using four swarms of eight particles each and PSO variables of w=0.73, c_1_= c_2_=1.5, and v_max_=1. The quality value reached a plateau value of 10809.0. On the other hand, the 3D model took 20.7 days using the same PSO setup. The cells in the basic model simulations accelerated for a relatively long time (300MCS) due the lack of the initial concentration field, therefore we added it. The cells then almost immediately jumped to the chemotaxis speed at the start of the simulation. Hence, we did not calculate the first 10 MCS steps, and obtained the correct speed with a small amount of minimization.

The optimized adhesion difference (*J*) values for each model, except the secreting and reference (1, 3, 4) ones, were not that particularly high compared to non-optimized ones. For example, 5_UB_ADH had (*J* of) 6.68, which is in the similar magnitude as 3_UB’s 5. However as expected, 8_ADH_ADH had the greatest variety of adhesion differences, e.g. in uniform model with: 0.50279 (‘NP and NP’) to 47.808 (‘Medium and NP’), and random model with: 0.50203 (‘NP and NP’) to 37.680 (‘MM and MM’) (Table 2). The chemotaxis lambda values were higher in the NP secreting models than UB secreting ones.

### Model Simulations

The correspondence of a simulation time unit (MCS) as an experimental one depended on the voxel (3D) scale (*l*), and the maximum speed of the relevant experiments (*v*_*max*_) (Andasari *et al*., 2012; Swat *et al*., 2012). And it can be calculated together with the physically maximum allowed speed (0.2 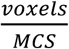) of the simulations

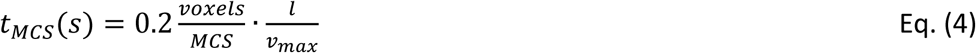

In our case, we supposed that a cell crawled on a substrate in an experiment with a speed of approximately 0.2 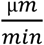. For the simulations, we assumed that 1 µm corresponds to 1 pixel in length. In other words, the lattice scale (*l*) was one 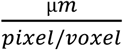. Then the maximum time per MCS was 0.2 * 1 / (0.2/60 [seconds/minute])) seconds = 60 seconds. Equation (4) allowed also the conversion of model speeds to same scope as experimental speeds.

Each simulation with a certain set of parameters and initial conditions was repeated 30 times to extract an average behaviour from the Monte-Carlo based algorithms. It was not possible to get simulation cell speeds to similar magnitude as observed experimentally merely by selecting the constants (of Eq. 2) arbitrarily, such as (*J*). Δ*H* depends also on local pixel concentration fields, and the other variables (Eq. 1-3). The chemoattractant flow is by default not hindered by the virtual cells, which is an approximation (Swat *et al*., 2012). There were no differences in the spread mechanisms of chemoattractant between (U and R) models with or without an initial chemoattractant field. CPM in the algorithm would try to modify all these, and then minimize the sum. According to Eq. (4), the maximum real time that can be assigned to 1000 MCS was ∼16.7h (Combes *et al*., 2016).

The simulated cell speeds, distances to corner and tip and amount of chemoattractant were derived from the changes of cell coordinates between each consecutive frame, more specifically from cell position changes in the centre of mass between each consecutive time step. Therefore, we used all time points for calculating these values. The chemoattractant concentration changes were not measured during the *ex vivo* and organoid experiments. We also calculated the ratio between tip and corner values of speeds and distances both in the simulations and experiments, should they be in different order of magnitude.

The ratios were needed because the corner distances of NP cells were within certain range. This constituted of the diameter of PTA with extra space. However, the moving (NP and MM) cells were in between the two UB trunks. At the same time, the tip distances of NP cells were in even smaller range in between the two tips. Hence, the ratio between tip and corner distances were needed to be also below one. Moreover, Combes experiment’s tip per corner NP cell speeds fluctuated relatively highly compared to model speed values. This was one of the reasons for giving separately corner speeds. The other reason was that, NP cells between or in the tips moved in swarming fashion.

### Technical Workflow

The emphasis of this study was to construct models and analyse computationally the resulting data. We wanted also to compare the model results to the experimental values of this and the previous (Combes et al.) work (Combes *et al*., 2016). However, the data frames obtained from simulations and experiments, were given in the form of ‘sample amount x time frame amount’, e.g. 30×1000 (for a model variant). Accordingly, the study design consisted of five separate phases, excluding making of the SOM and PSO routines, described elsewhere in the appendix.

Shortly; importing data to a coding platform (SPYDER), unifying data frames of experiments and models (.CSV with PANDA), sample selection (NP or MM cells from the regions), calculations of speeds, distances and concentrations with specific functions (i.e. the mean in each time point and cell across all the samples), and expressing the averages or plots of the previous measures.

Following importation of model and experimental data into a coding development platform (SPYDER, using python (Contributors, 2017)) the data frames were unified (using the PANDA package (Contributors, 2017)), symmetrised and converted to similar between model and experimental samples. The relevant experimental samples were selected for calculations. The corresponding NP and MM cells assigned closest to the left UB tip in simulations were selected for calculations (Fig. 8A). Finally, the average values for these samples and cells were calculated and values plotted.

In their original analysis Combes et al. (Combes *et al*., 2016) considered only NP cell coordinates whereas we simulated the behaviour of both NP and MM cells between two adjacent UB tips. Presumably also in this spatiotemporal setting the cells surrounding either UB tip reacted first to the chemoattractant coming from that UB (Short *et al*., 2014), as they did in other previous experiments (Lefevre *et al*., 2017). Presumably, this applied also to the chemoattractant coming from the secreting NP cells closer to each other (Fig. 2D/F). This practically meant that we give the average measures for those NP and MM cells closer to left-hand side L-structure.

Consequently, the calculations of speeds, distances and concentrations were performed for NP and MM cells near to the left-hand side of the simulated UB (c.f., Figs 1D/E, 8A/E). Therefore, the experimental and simulated distances of cells to the tip or corner were comparable. This was mainly a result of similar spatial dimensions and effective cell motilities in simulated vs. experimental systems. The experimental tip and corner distances were normalized (0 to 1), followed by multiplying the transformed distances by model averages. This was done because the frame shapes in the experiments and simulations were slightly different (Figs 1, 8). The speeds and concentrations were not scaled between simulations and models. However, we give the tip per corner values for the cell distances and concentrations. This was because we needed to scale the experimental (tip and corner) distances to the same order of magnitude as the model values.

In addition, during the work of setting the model, we discovered that the value of the chemotaxis secretion constant multiplied by its strength constant, i.e. the chemotaxis lambda, should be less than twenty. The reason for this was to avoid a virtual cell loss. Otherwise, the loss would arrive from the disruption of the virtual cell membranes. It was possible that same would apply in the early nephrogenesis, i.e. cells would need to control the amount of secretion and how to respond to it. Therefore, the average dimensionless concentration levels were handy way to assess the reliability of the workflow for a stable model simulation (Figs 8, S2, S6, S7). This did not of course apply to the non-secreting models (1, 2, 8), which did not have any chemoattractant (Fig. 2).

### Details of SOM

SOM is a machine learning method involving an artificial network to represent the original experimental or simulation data. The method resembles mathematical iteration by conducting an unsupervised learning to compress the data to a two-dimensional (2D) map of nodes. The primary relationships of the data elements, such as the cell names and their different coordinates, can be conserved with a neighbourhood function. (Kohonen, 1982)

The iteration routine consisted of giving weights to the real distances between the network nodes and the data. The node whose weight vector was most similar to the data was called the best matching unit (BMU). The weights of the BMU and nodes close to it in the SOM grid were adjusted towards the input vector. The magnitude of the change decreased with time and with the grid-distance from the BMU. The iteration equation for a node *v* with weight vector *W*_*v*_ was

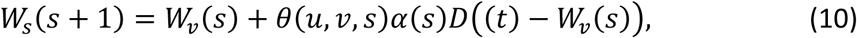

where *D(t)* was the data vector, *s* was the iteration step index number, *t* was a randomly or otherwise selected index number of the training sample, *u* was the index number of the BMU for *D(t)*, and *α(s)* was a monotonically decreasing learning coefficient. Finally, *θ(u,v,s)* was the neighbourhood function. It gave a distance between the previous node (*u)* and the node (*v)* in the iteration step (*s)*. (Kohonen, 1982)

For performing this kind of an iterative calculation, the use of computer program techniques was naturally implied, such as ‘MiniSom’ python function developed earlier (GitHub, 2019). Shortly, it required four instructions: 1. Importing and normalizing the initial experimental or simulation data, 2. Specifications for the node map; the size of the node map matrix, learning coefficient, initial spread of neighborhood function as gaussian, and the neighborhood function, which was here the ‘Mexican hat’ in order to gather all 2D nodes inside an oval, 3. By randomly and with the principle component analysis style weighted BMUs, and 4. The iteration, i.e. the random training 5000 times by default, since more than 300 did not enhance the accuracy significantly (GitHub, 2019), and plotting the SOM nodes with gradually increasing colors in respect to their closeness to the original data. As for the other values, sigma was four, learning rate 0.5, and the size of the random seed 10. The routine took about 15-40 minutes depending on the original size, i.e. memory space used by the computer, of the initial data. This was followed by the grouping of closest SOM nodes, or groups, to different categories, such as here to the cell coordinates and their instantaneous speeds, in respect to the previous time point coordinates.

The segmentation worked so that we reduced the noise-cell coordinates to their respective real cells by applying the mean area and speed (50 µm^2^, 0.15 µm/min) (Stegmaier *et al*., 2016). The tool selected and grouped all the cell coordinates together with their speeds in the experiments or simulations. With the appropriate SOM transformed plotting routines, we evaluated the reasons for the cell developments in the different cell coordinates.

It was not necessary to find the NP cell speed regions with SOM or otherwise by choosing certain NP cells rather than the derivates of their speed trend. This was performed by selecting all instantaneous NP speeds and their coordinates of experimental noisy data (particularly our experiments) to SOM. And after the iteration procedures, we transformed the speed contours, i.e. ‘stable regions’, back to real normed space with experimental images in the background (Stegmaier *et al*., 2016). As a result, SOM could be also used as a way to check the imaging or segmentation accuracy, as well as a segmentation tool as such.

We devised a SOM node matrix size as 10 times 10 due computational efficiency (GitHub, 2019), and the original size of the experimental data matrix as 25000 times four (Combes *et al*., 2016; GitHub, 2019). And from those 100 possible nodes (of SOM), we found 10-30 nodes in total that were close to the original data (after the iterative training). Subsequently, we chose eight closest groups to the data.

We further selected the groups, where the cell speeds were 95 % similar in the beginning and in the end. This was because we were interested in the different regions of speeds. There were usually three groups from the eight. We could remap these three nodes of ‘speeds and coordinates of NP cells’ back to normalized 2D or 3D experimental or simulation space, because we knew the (original) coordinates of these cells.

## Captions of Supplementary Images

**Fig S1.**
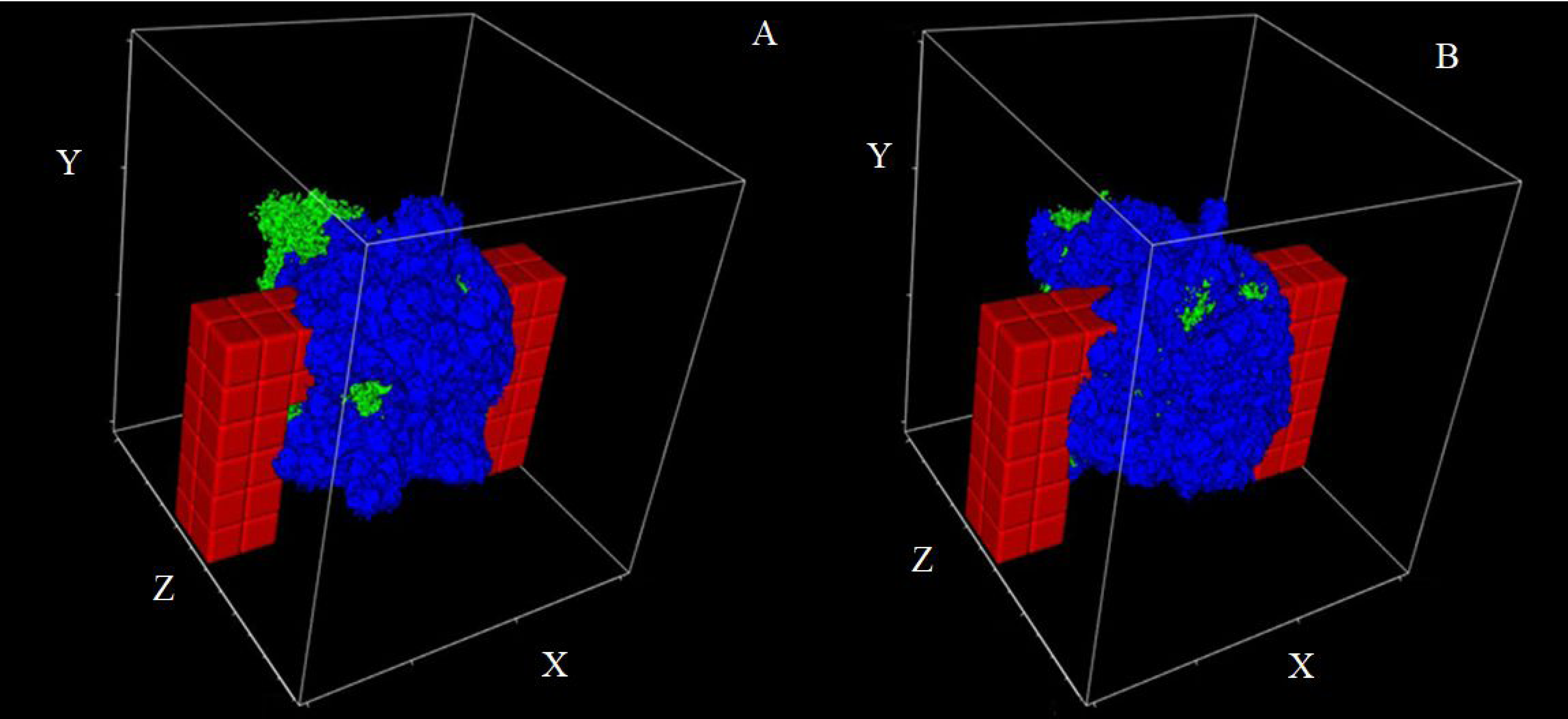
Cell patterns at the end of CC3D simulations. The NP secreting model (4_NP) was simulated with the initially A) uniform and B) random cell patterns (defined in Fig. 8). UB cells are depicted with red, NP cells with green, and MM cells with blue colours.

**Fig. S2.**
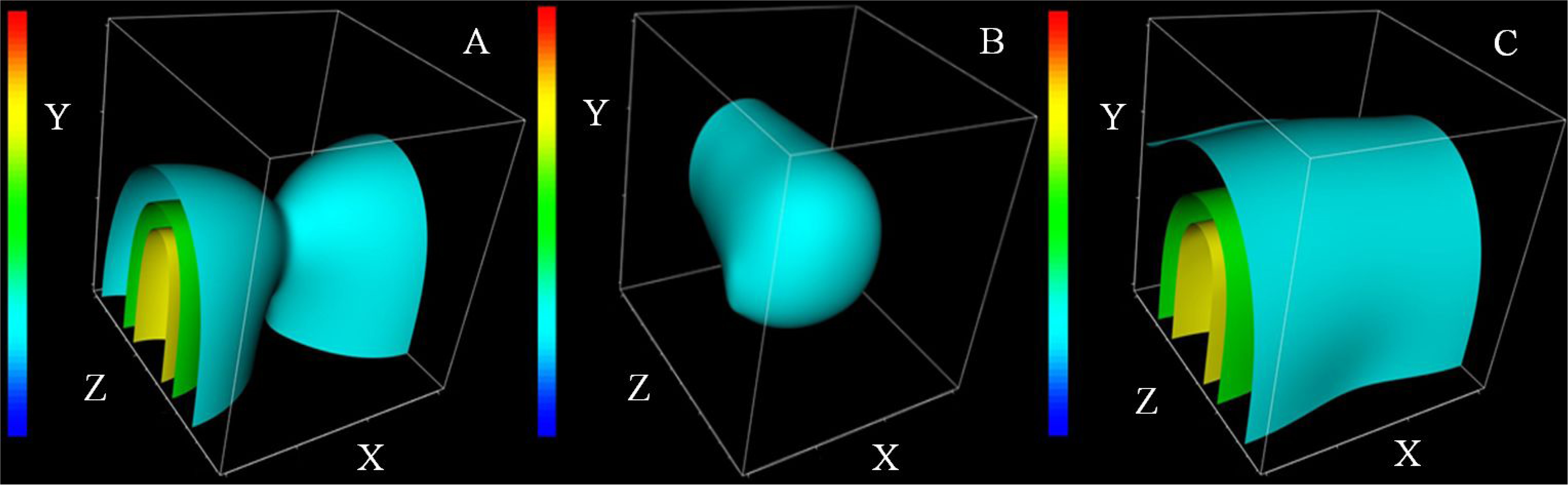
Chemoattractant patterns at the end of CC3D simulations. The primary patterns for UB and NP secreting models. A) 3_UB and 5_UB_ADH, B) 4_NP and 6_NP _ADH, C) 7_UP_NP_ADH. Standardized chemoattractant concentration gradients are depicted by sheets, ranging from 0 (blue) to 1 (red).

**Fig. S3.**
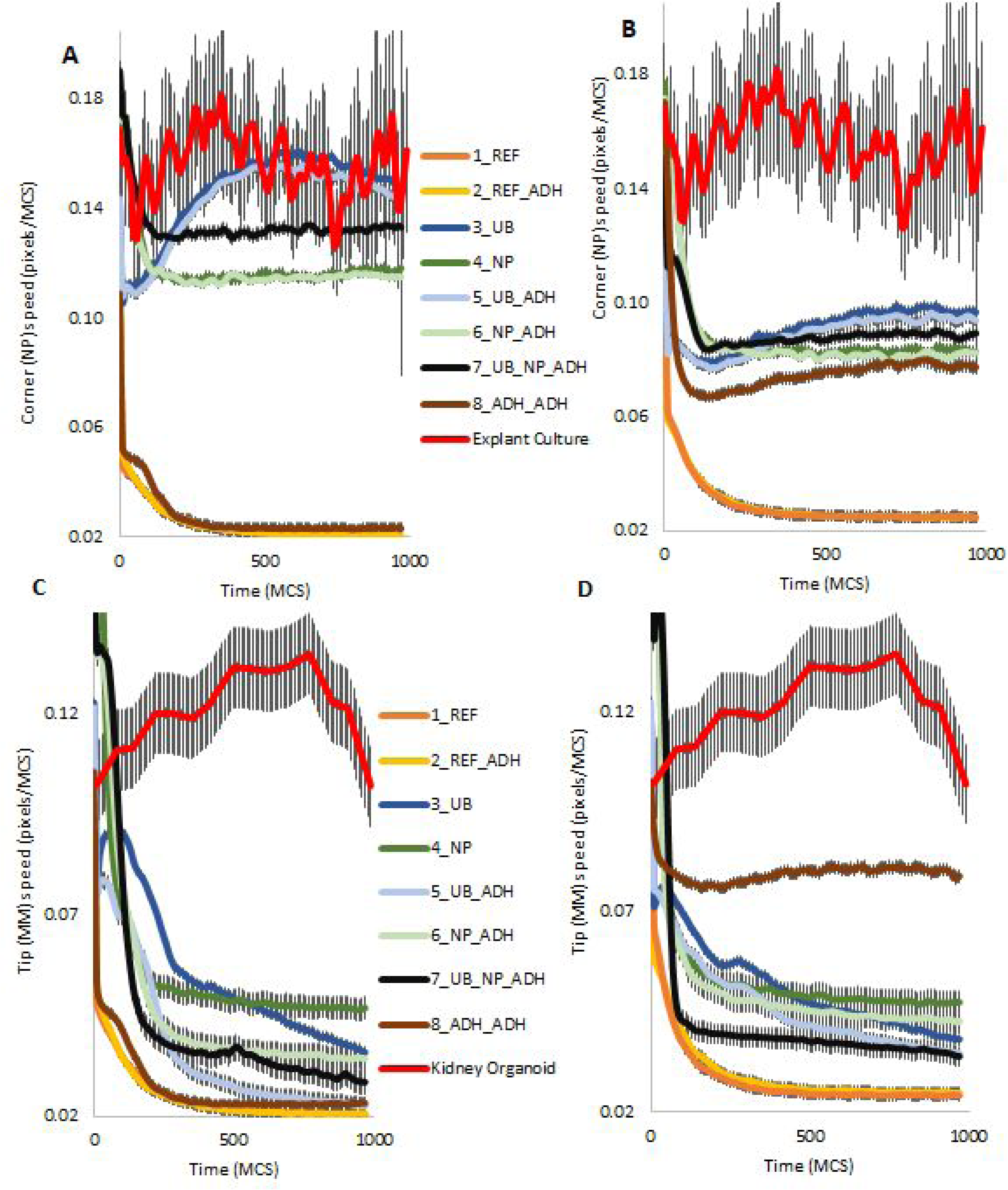
Average speeds of NP cells in tip and MM cells in corner in the simulation studies of different models together with the results from experiments. A/B) NP cells in the models (see legend) with the cells in the explant culture experiments (Combes *et al*., 2016). C/D) MM cells in the models (see legend) with cells in the kidney organoid experiments. A/C) Simulations before the optimization. B/D) Simulations after the optimization. Vertical bars indicate 95% confidence intervals.

**Fig. S4.**
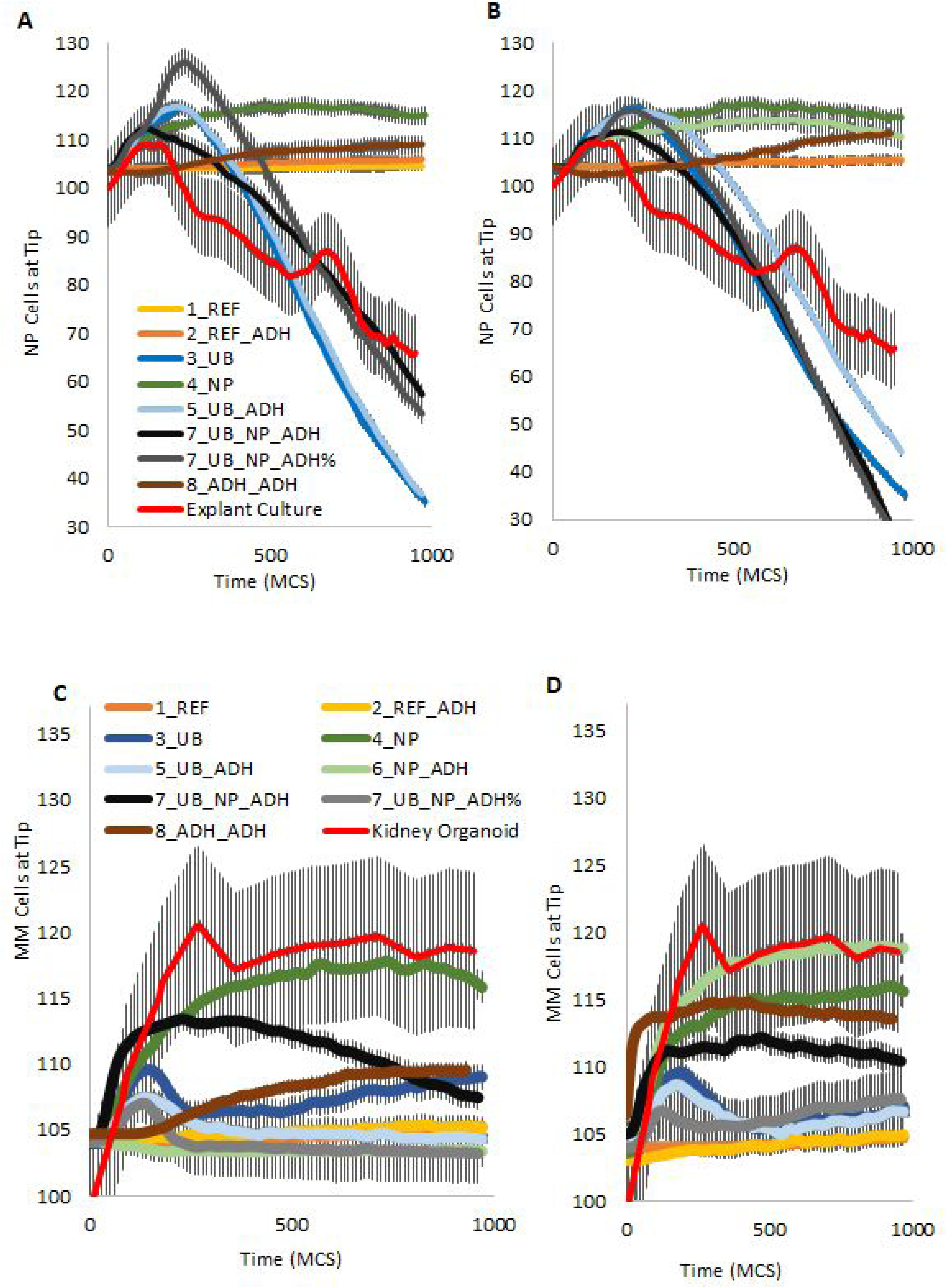
Average NP (A/B) and MM (C/D) cell quantities at the tip in the simulation studies of different models together with the results from experiments. A/B) NP cells in the models (see legend) with the cells in the explant culture experiments (scaled) (Combes *et al*., 2016). C/D) MM cells in the models (see legend) with the cells in the kidney organoid experiments (scaled). A/C) Simulations before the optimization. B/D) Simulations after the optimization. Vertical bars indicate 95% confidence intervals.

**Fig. S5.**
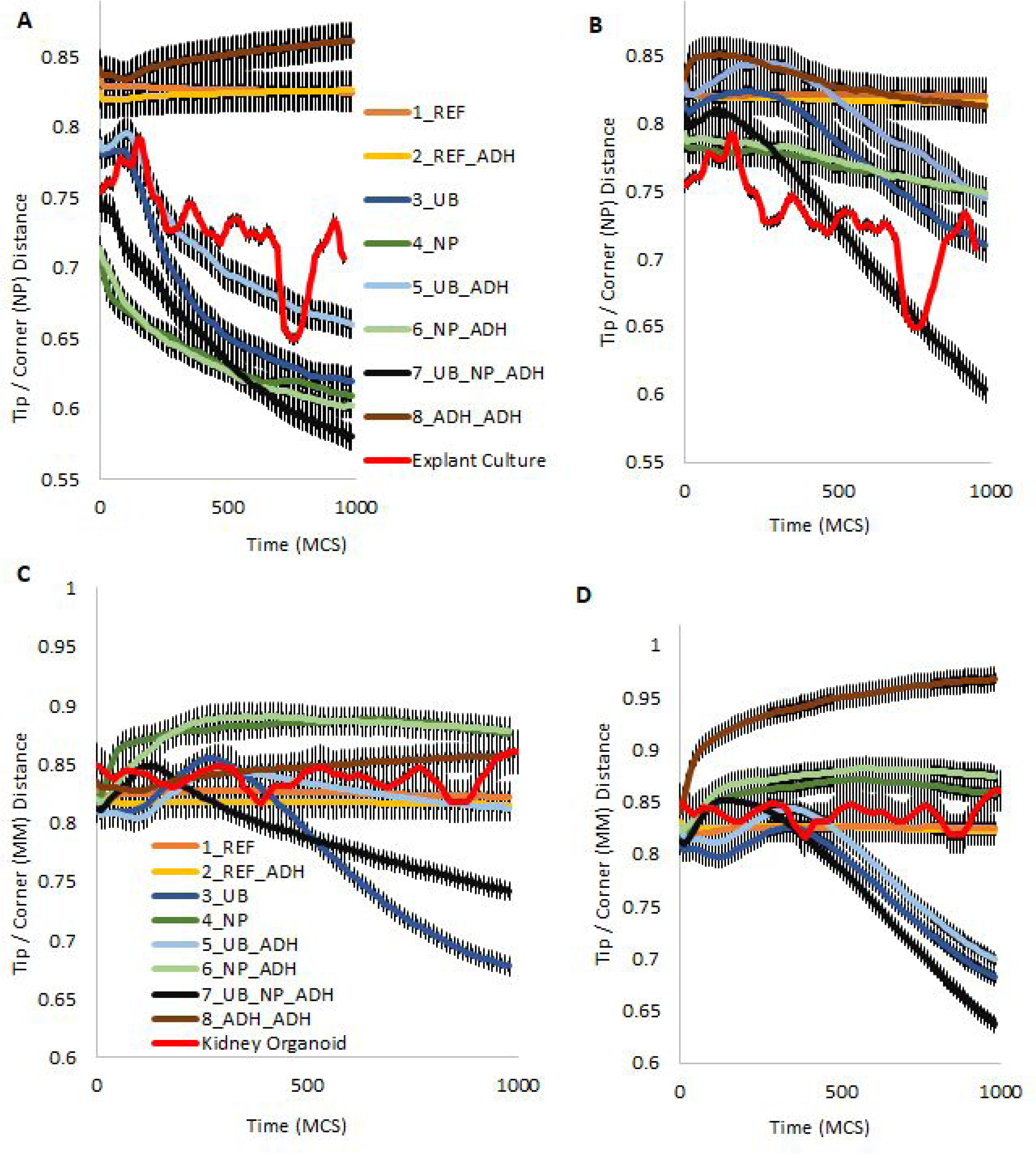
Tip-to-corner distance ratios of NP (A/B) and MM (C/D) cells in the simulation studies of different random models together with the results from experiments. A/B) NP cells in the models (see legend) with the cells in the explant culture experiments (scaled to model 3 and 4) (Combes *et al*., 2016). C/D) MM cells in the models similarly scaled with cells in the kidney organoid experiments. A/C) Simulations before the optimization. B/D) Simulations after the optimization. Vertical bars indicate 95% confidence intervals.

**Fig. S6.**
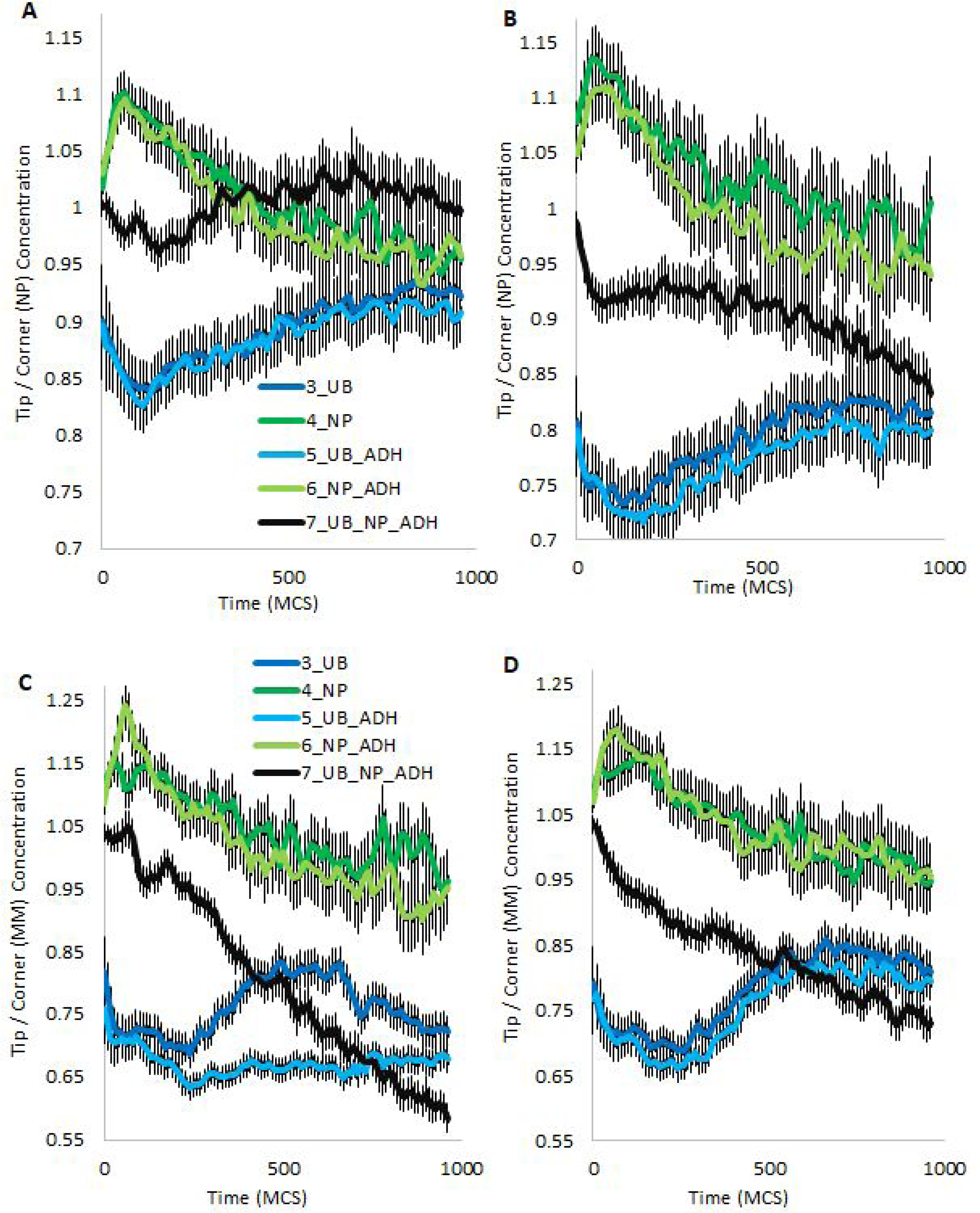
Average tip per corner concentration changes in COMs of NP and MM cells in the simulation studies of different models together with the results from experiments. A/B) NP cells in the models (see legend) with the cells. C/D) MM cells in the models. A/C) Simulations before the optimization. B/D) Simulations after the optimization. Vertical bars indicate 95% confidence intervals.

**Fig. S7.**
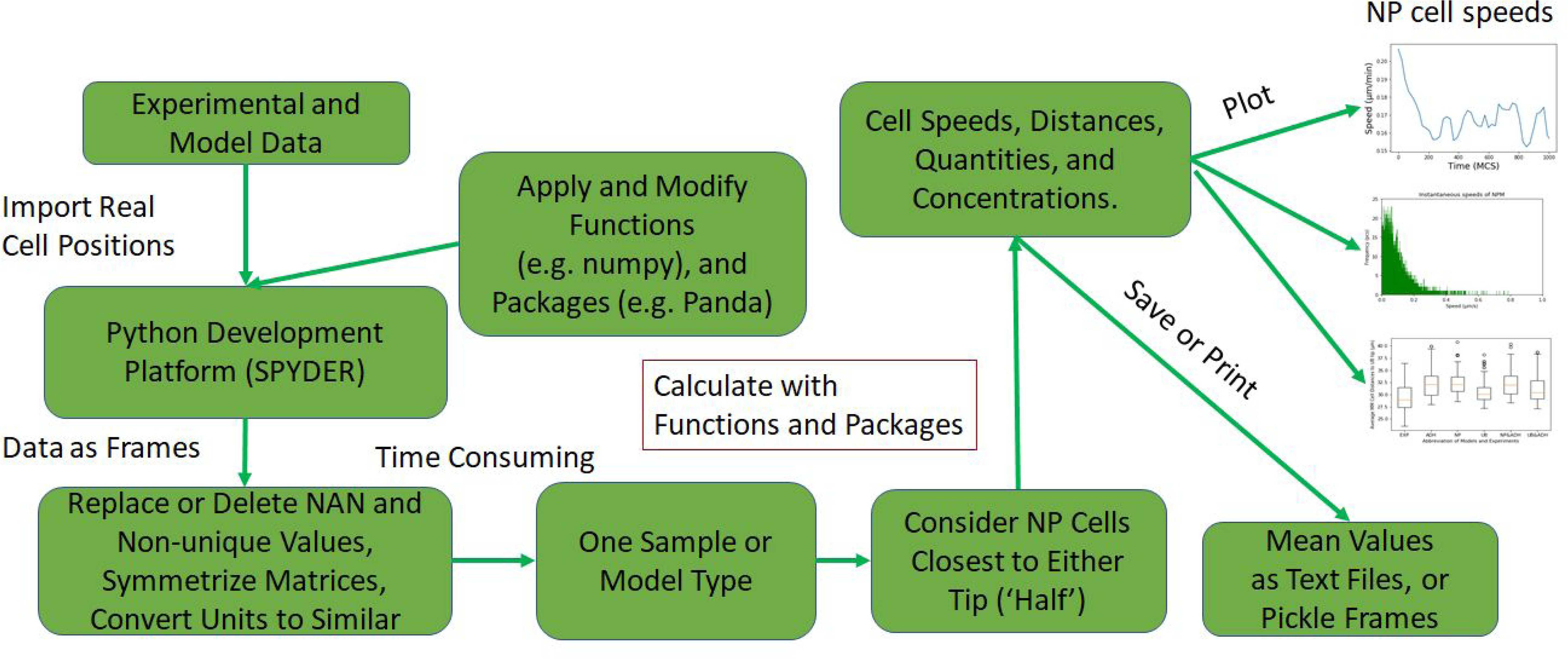
Study design of computational analysis workflow. Experimental and model data was importer to Spyder and converted to symmetrical matrices. The calculation of NP cell speeds and positions were done after selecting matching samples of experiments and models, and plotted with own functions.

